# A Quantitative Sequencing Framework for Absolute Abundance Measurements of Mucosal and Lumenal Microbial Communities

**DOI:** 10.1101/2020.02.28.970087

**Authors:** Jacob T. Barlow, Said R. Bogatyrev, Rustem F. Ismagilov

## Abstract

A fundamental goal in microbiome studies is to determine which microbes affect host physiology. Standard methods for determining changes in microbial taxa measure relative microbial abundances, which cannot capture absolute changes. Moreover, studies often focus on a single site (usually stool), although microbial demographics differ substantially among gastrointestinal (GI) locations. Here, we developed a quantitative framework to accurately measure absolute abundances of individual bacterial taxa by combining the precision of digital PCR with the high-throughput nature of 16S rRNA gene amplicon sequencing. In a murine ketogenic-diet study, we compared microbial loads in lumenal and mucosal samples at several sites along the GI tract. Measurements of absolute (but not relative) abundances revealed decreases in total microbial loads on the ketogenic diet and enabled us to accurately determine the effect of the diet on each taxon at each GI location. Quantitative measurements also revealed different patterns in how the ketogenic diet affected each taxon’s abundance in stool and small-intestine mucosa samples. This rigorous quantitative microbial analysis framework applied to samples from relevant GI locations will enable mapping microbial biogeography of the mammalian GI tract and more accurately capture the changes of microbial taxa in experimental microbiome studies.

---

One main goal of microbiome studies is to determine which taxa, if any, drive phenotypic changes among study groups.^1-3^ The first step in this process is often to survey which microbial taxa differ in abundance between study groups (differentially abundant taxa). This survey is commonly performed by amplifying the 16S rRNA gene amplicon with “universal” primer sets before high throughput sequencing.^4^ The output of these studies provides the relative, not absolute, abundance of each taxon in each sample. Researchers often then use standard statistical tests or microbiome specific packages to determine which taxa are differentially abundant.^5, 6^

Relative-abundance analyses are effective for determining the major microbial taxa in an environment (e.g., the human Microbiome Project). However, several researchers have pointed out the inherent limitations of comparing relative abundances between samples.^7-10^ In analyses of relative data, every increase in one taxon’s abundance causes an equivalent decrease across the remaining taxa. Thus, the measurement of a taxon’s relative abundance is dependent on the abundance of all other taxa, which can lead to high false positive rates in differential taxon analyses^8, 11-13^ and negative-correlation biases in correlation-based analyses.^14, 15^ Several methods (e.g., ALDEx2^16^, Ancom^17^, Gneiss^18^, Differential Ranking^10^) acknowledge these biases and aim to address them by using the ratios among taxa, which are conserved regardless of whether the data are relative or absolute. These methods are particularly valuable because they enable improved re-analysis of existing datasets reporting relative abundances.^10, 16-18^

Despite such methodological advancements, analyses of relative abundance cannot fully capture how individual microbial taxa differ among samples or experimental conditions. Using the simple example of a community containing two taxa (Fig. 1), we see that an increase in the ratio between Taxon A and Taxon B could indicate one of five scenarios: (i) Taxon A increased (Fig. 1a), (ii) Taxon B decreased (Fig. 1b), (iii) A combination of 1 and 2, (iv) Taxon A and Taxon B increased but Taxon A increased by a greater magnitude, or (v) Taxon A and Taxon B decreased but Taxon B decreased by a greater magnitude (Fig. 1c). Knowing which of these five scenarios occurs when analyzing experimental data could drastically alter the interpretation of which taxa are positively or negatively associated with phenotypes. Thus, an inherent limitation of methods that use relative abundance is that they cannot determine whether an individual taxon is more abundant or less abundant (the direction of the change) or by how much (the magnitude of the change) between two experimental conditions or samples.

**Figure 1:**
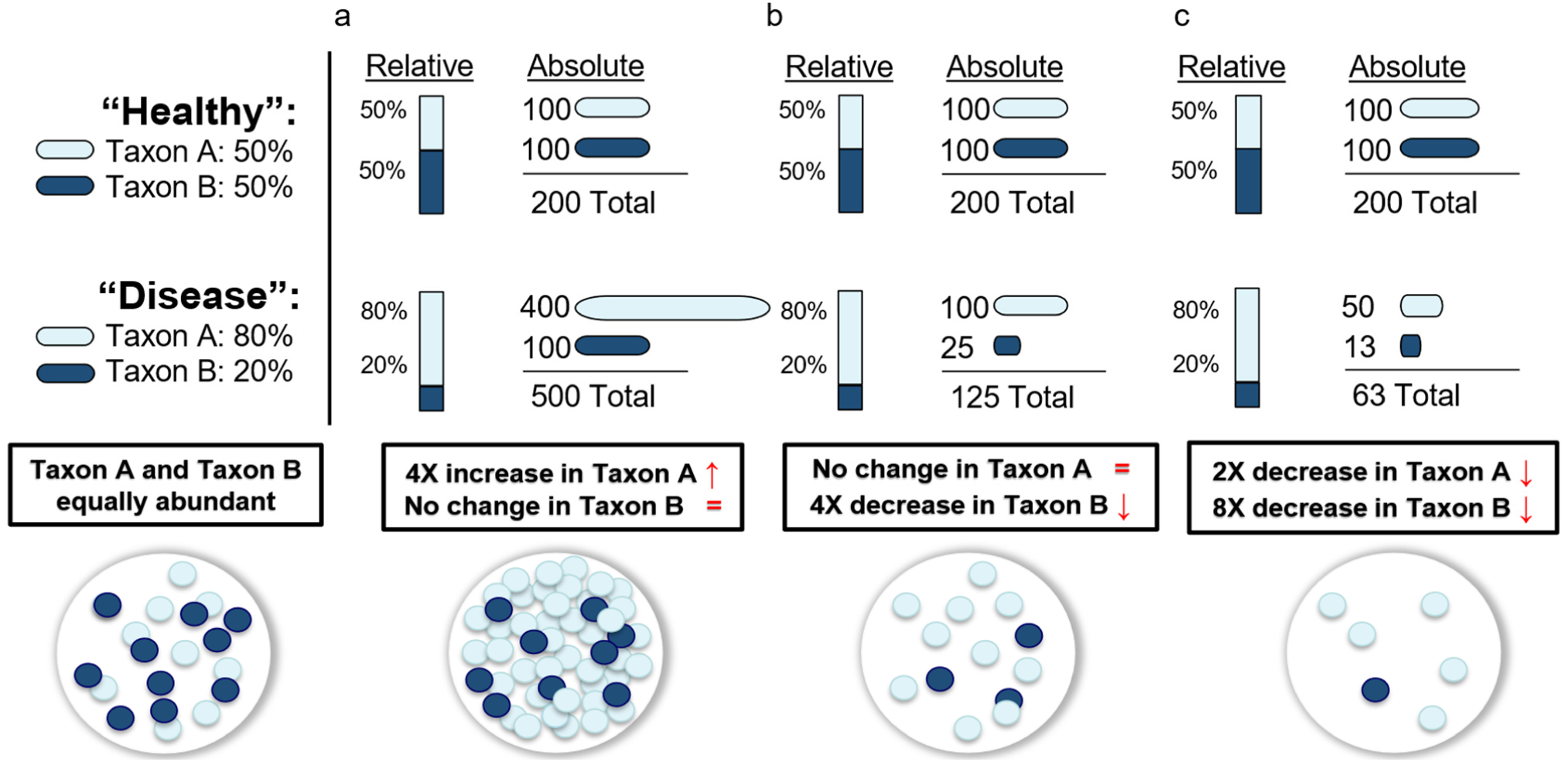
The value of absolute quantification is illustrated by three hypothetical scenarios in which the relative abundance of two taxa (Taxon A and Taxon B) are found in equal abundance (50:50) in a “healthy” state but in an 80:20 ratio in the “disease” state. (a) Taxon A increases in abundance while Taxon B remains the same; (b) Taxon A remains unchanged while Taxon B decreases in abundance, and (c) Taxon A and Taxon B both decrease, but Taxon B decreases by a greater magnitude.

To overcome these limitations, several important methods have been developed for quantifying the absolute abundance of microbial taxa by using known “anchor” points to convert relative data to absolutes. Spiked standards are commonly used in method calibration and have recently been applied to quantifying taxa in microbiome research.^19-23^ These methods require a purified DNA sequence of known concentration from an organism not present in the sample and an estimate of the initial sample concentration to determine the amount of exogenous DNA to spike-in. Another group of anchoring methods, such as those that use flow cytometry^24^, total DNA^25^, or qPCR^26-28^, measure the total concentration of cells, DNA, or amplicons to transform the relative abundances to absolute numbers. These methods have already demonstrated the value of quantitative microbiome analysis, yet microbiome researchers have not yet uniformly adapted these methods. One may speculate that this lack of adoption is because of real or potential limitations of these methods. For example, flow-cytometry based methods require dissociating the sample into single bacterial cells, which could require complex sample preparation and have not been validated with complex samples such as from gut mucosa. Total-DNA-based methods are limited to samples only containing microbial DNA (no host DNA), and spike-in or qPCR-based methods can be limited by amplification biases.^29, 30^ To increase utilization of quantitative microbiome analyses, the following capabilities and validation need to be demonstrated: (i) performance across samples with microbial loads ranging from high, as in stool, to low, as in the small intestine; (ii) performance across biogeographically diverse sample types, from microbe-rich stool and colonic contents to host-rich mucosal samples; (iii) explicit evaluation of limits of quantification of the method, and how these limits depend on the starting microbial load, relative abundance of a specific target taxon in the sample, and sequencing depth.

To address this challenge, in this paper we establish a rigorous, absolute quantification framework based on digital PCR (dPCR) anchoring. We chose dPCR as our anchoring method because PCR is already part of sequencing protocols and has been extensively validated as a quantitative method in nucleic-acid measurements. To achieve precise measurements of absolute abundance from diverse sample types, we assessed the efficiency and evenness of the DNA extraction protocol. To minimize and quantify bias resulting from potentially uneven amplification of microbial 16S rRNA gene DNA, or non-specific amplification of host DNA, we utilized dPCR in a microfluidic format.^31-33^. dPCR is an ultrasensitive method for counting single molecules of DNA or RNA.^34-36^ By dividing a PCR reaction into thousands of nanoliter droplets and counting the number of “positive” wells (those with amplified template), dPCR yields absolute quantification without a standard curve. To understand the quantitative limits of our methodology, we measured the accuracy of each taxon’s absolute abundance as a factor of both input DNA amount and individual taxon relative abundance.^37-39^ We then evaluated this absolute quantification workflow by performing a murine ketogenic-diet study that illustrates how the selection of relative-vs. absolute-quantification analyses can result in different interpretations of the same experimental results. Many studies have shown that ketogenic diets can induce substantial compositional changes in gut microbiota,^40-42^ so, we predicted it would serve as a good illustrative model for our workflow. Finally, we applied this workflow to an analysis of microbial loads along the entire gastrointestinal (GI) tract to highlight the importance of judicious selection of sample location when evaluating the impact of diet on host phenotype, and to highlight the applicability of this workflow to GI sites with diverse microbial loads.

## Results

### Microbial DNA extraction was efficient, unbiased, and quantitative over a wide range of microbial loads and across intestinal sample types

To estimate the maximum quantity of sample we could extract before overloading the 20-µg column capacity, we measured total DNA and microbial DNA load across small intestine and large intestine lumenal and mucosal samples (Fig. S1). We then evaluated extraction efficiency across three tissue matrices (mucosa, cecum contents, and stool) to assess whether variation in levels of PCR inhibitors and non-microbial DNA interfered with microbial quantification. We spiked a defined 8-member microbial community into GI samples taken from germ-free (GF) mice. To assess quantitative limits, we performed a dilution series of microbial spike-in from 1.4 × 10^9^ CFU/mL to 1.4 × 10^5^ CFU/mL. dPCR quantification showed near equal and complete recovery of microbial DNA over 5 orders of magnitude (Fig. 2a). Overall, we measured ∼ 2X accuracy in extraction across all tissue types (cecum contents, stool, SI mucosa) when total 16S rRNA gene input was greater than 8.3 × 10^4^ copies (Fig. S2). Normalizing this sample input to the approximate maximum extraction mass (200 mg stool, 8 mg mucosa) yielded a lower limit of quantification (LLOQ) of 4.2 × 10^5^ 16S rRNA gene copies per gram for stool/cecum contents and 1 × 10^7^ 16S rRNA gene copies per gram for mucosa. Mucosal samples had a higher LLOQ because the high host DNA in this tissue type saturates the column, limiting total mass input.

**Figure 2:**
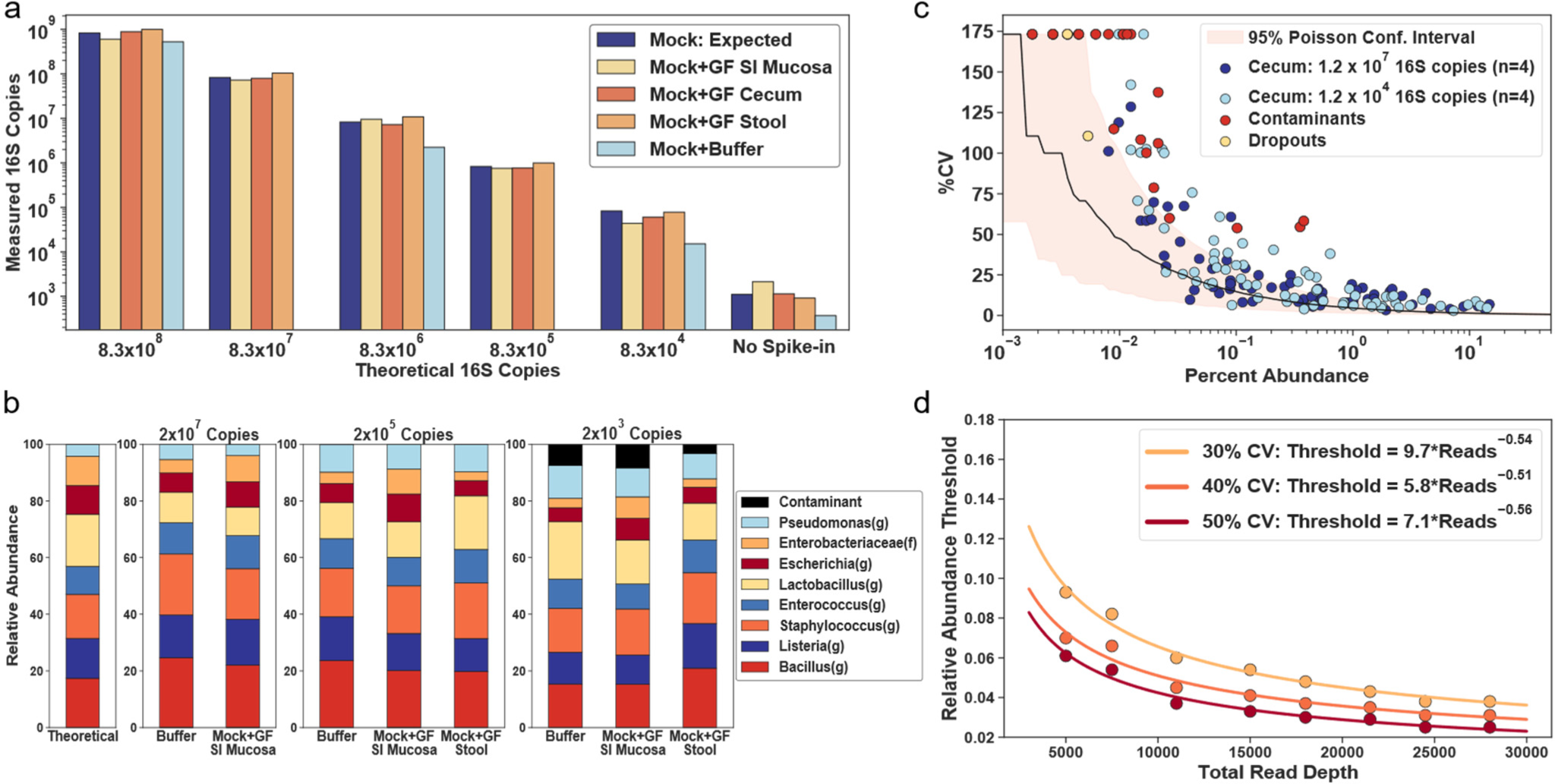
Lower limits of quantification for total microbial DNA extraction and 16S rRNA gene amplicon sequencing. (a) A comparison of theoretical and measured copies of the 16S rRNA gene with digital PCR using an eight-member microbial community spiked at a range of dilutions into germ-free (GF) mouse tissue from small-intestine (SI) mucosa, cecum, and stool. Each bar plot shows a single technical replicate for each matrix. (b) Relative abundance of the eight taxa as predicted and measured after 16S rRNA gene amplicon sequencing. (c) Correlation between the mean (n=4) relative abundance of each taxon and the coefficient of variation (%CV) using a cecum sample from a mouse on a chow diet with an initial template input of either 1.2 × 10^7^ or 1.2 × 10^4^ 16S rRNA gene copies. Each analysis comprised four technical (sequencing) replicates. Taxa found only in the low-input sample were labeled contaminants (red points); taxa found in the high-input sample but not low input sample were labeled dropouts (yellow points). Red shading indicates the Poisson sampling confidence interval (10,000 bootstrapped replicates) at a sequencing read depth of 28,000. (d) Relationship between relative abundance threshold (see text for details) and sequencing read depths at 30%, 40%, and 50% CV thresholds.

Next, to ensure extraction performance was consistent for both Gram-negative and Gram-positive microbes, we performed 16S rRNA gene amplicon sequencing using previously described improved primers and protocol^31, 33^ on a subset of the extracted samples (Fig. 2b). It is important to note that all amplification reactions for 16S rRNA gene library prep were monitored with real-time qPCR and we stopped the reactions when they reached the late exponential phase to limit overamplification and chimera formation.^30–33, 43, 44^ Extraction appeared less even among microbial taxa at lower total microbial DNA inputs (Fig. 2b). This discrepancy from the theoretical profile did not correlate with the presence of chimeric sequences (Fig. S3) and was likely a function of the reduced accuracy incurred when diluting complex microbial samples. Additionally, sequencing samples with low total microbial loads (<1 × 10^4^ 16S rRNA gene copies) resulted in the presence of contaminants, as confirmed by sequencing of negative-control extractions (Table S1).

### Quantitative limits of amplicon sequencing provide informative thresholds for data analysis

To establish the precision of relative-abundance measurements, we sequenced four replicates of DNA extractions from cecum samples. Libraries from one DNA extraction were prepared with either an input of 1.2 × 10^7^ 16S rRNA gene copies or 1.2 × 10^4^ 16S rRNA gene copies to determine the impact of starting DNA amount on sequencing variability. We calculated the coefficient of variation (%CV) for each taxon’s relative abundance from amplicon sequencing the replicate samples. Each taxon’s mean relative abundance (n=4) was then plotted against its corresponding coefficient of variation of the relative abundance (Fig. 2c). We defined “dropouts” as taxa present only in the high-DNA-input sample whereas we defined “contaminants” as taxa present only in the low-DNA-input sample. The two dropout taxa in the low input sample corresponded to the lowest abundance taxa from the high input DNA sample (yellow points, Fig. 2c). Most of the contaminant taxa had a relative abundance < 0.03%, but three taxa (*Pseudomonas(g), Acinetobacter(g), Rhizobiales(f)*) had relative abundances of 0.38%, 0.35%, and 0.1%, respectively. These three taxa were also the three most common contaminants in our negative-control extractions (Table S1). The presence of contaminants in the sample containing 1.4 × 10^4^ 16S rRNA gene copies was consistent with the input amount at which we observed contaminants in our mixed microbial community dilutions (Fig. 2b). We calculated a bootstrapped Poisson sampling confidence interval at our sequencing depth (28,000 reads) to assess how close our accuracy limits were to the theoretical limits (red shading, Fig. 2c). At the low DNA input level of 1.2 × 10^4^ 16S rRNA gene copies, we began to reach the fundamental Poisson loading limit in our library-preparation reaction (Fig. S4a). We expected divergence of the %CV at ∼0.01% abundance because at a read depth of 28,000 a relative abundance of 0.01% is a measure of ∼3 reads whereas at a total 16S rRNA gene copy input of 1.4 × 10^4^ a relative abundance of 0.01% is ∼1 copy. Poisson statistics also helped us define the theoretical lower limits of relative-abundance measurements as a factor of sequencing depth (Fig S4b).

We next wished to quantify an approximate threshold that would tell us, for a given sequencing depth, at what percentage of relative abundance we lose accuracy in our measurements (we defined this threshold as “relative abundance threshold”). To determine this threshold, we fit a negative exponential to the replicate data and identified the percentage abundance at which 30% CV was observed. This threshold is a function of the sequencing depth, so we subsampled the data at decreasing read counts and repeated the exponential fitting method to calculate the relationship between the relative abundance threshold and sequencing depth (Fig. 2d). Greater sequencing depths yielded lower quantitative limits with diminishing returns, as expected. We found that the threshold for percentage abundance decreases with increasing sequencing depth with a square root dependence analogous to the square-root dependence of Poisson noise. This trend follows for %CV thresholds of 40% and 50% as well (Fig. 2d). This analysis provides a framework with which to impose thresholds on relative-abundance data that are grounded on the calculated limits of quantitation.

### Digital PCR anchoring quantifies bias in amplicon sequencing and provides a framework for absolute quantification of taxa

We calculated absolute abundances of taxa from sequencing data using dPCR measurement of total microbial loads as an anchor. Briefly, relative abundance of each taxon was measured by sequencing and these numbers were multiplied by the total number of 16S rRNA gene copies (obtained using the same universal primers from amplicon sequencing) from dPCR. Next, we evaluated the accuracy of this quantitative sequencing approach. We were not able to directly compare our measurements to other absolute abundance techniques discussed in the introduction because these techniques have not been validated on the broad range of sample types and microbial loads tested here (Table S2). A fair side-by-side comparison would require re-optimization of current techniques for complex sample types like those with high host DNA levels and low microbial biomass (e.g., mucosa). Typically, evaluation of quantitative accuracy and precision would involve the use of a mock microbial community (like the one used in Fig. 2). However, because we computed the absolute instead of relative abundances, we were able to use the actual gut-microbiota samples and compare the results to the dPCR data obtained with relevant taxa-specific primers. The 16S rRNA gene copy amount was then normalized to the mass of each extracted sample after correcting for volume losses (Materials and methods; Eqn. 1). We chose four representative taxa to encompass common gut flora of varying classification levels: *Akkermansia muciniphila(s), Lachnospiraceae(f), Bacteroidales(o)*, and *Lactobacillaceae(f)*. Like eubacterial primers, taxa-specific primer sets can (in principle) give rise to nonspecific amplification due to overlap with host mitochondrial DNA. To avoid nonspecific amplification, we ran temperature gradients with GF mucosal DNA and taxa-specific microbial DNA to identify the optimal annealing temperature for each primer set (Fig. S5). Each taxa-specific primer targets a separate region of the 16S rRNA gene than the universal primer set, thus keeping the gene copy number equivalent across primers. We observed high correlation coefficients between the taxa load determined by quantitative sequencing with dPCR anchoring and the taxa load measured by dPCR with taxa-specific primers (all r^2^ >= 0.97, Fig. 3a) for all four taxa over a range of ∼ 6 orders of magnitude. The ratio of the total load measurements obtained by quantitative sequencing with dPCR anchoring and by dPCR with taxa-specific primers showed unity agreement between three of the four primer sets with 2-fold deviation from the mean (Fig. 3b and Fig. S9). Sequencing quantification was consistently 2.5-fold higher than dPCR quantification for the species *Akkermansia muciniphila* (Fig. 3b). We cannot confirm amplification bias as a factor because the error did not depend on the number of cycles used in library preparation. An alternative factor could be a discrepancy in coverage/specificity between the taxon-specific and universal primer sets. We next tested the limits of the sequencing accuracy as a factor of input DNA load. A 10X dilution series of a cecum sample was created to cover input DNA loads of 1×10^8^ copies down to 1×10^4^ copies. Minimal differences in beta diversity (Aitchison distance) between the undiluted and diluted samples were observed with a trend towards increasing difference with decreasing DNA load (Fig. 3c). This negative correlation between beta diversity and microbial load is not unexpected due to the higher presence of contaminant species from our negative controls in the lower input samples, specifically *Pseudomonas*(g) (data not shown).

**Figure 3:**
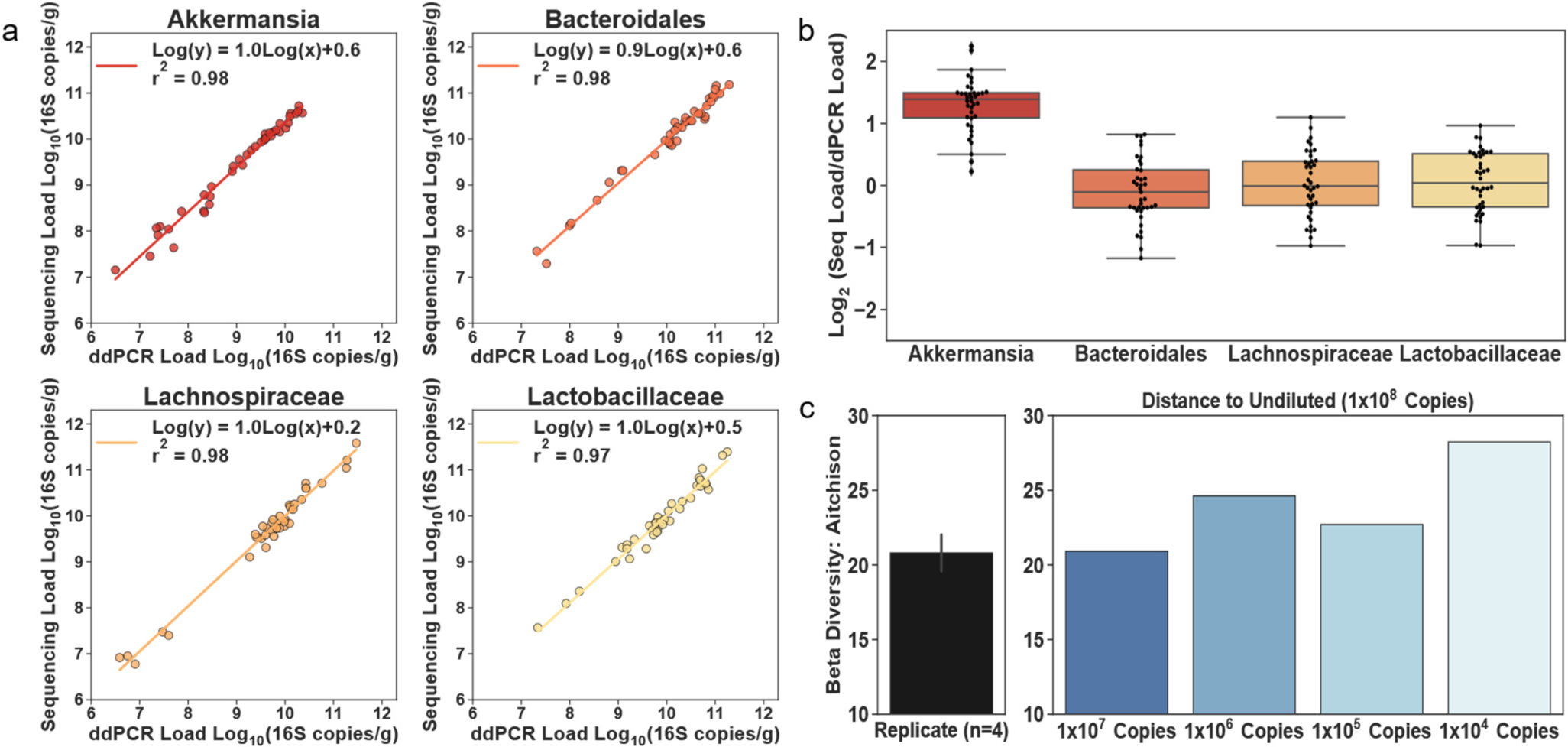
Taxon-specific digital PCR (dPCR) demonstrates low biases in abundance measurements calculated by 16S rRNA gene quantitative sequencing with dPCR anchoring. (a) Correlation between the Log_10_ abundance of four bacterial taxa as determined by taxa-specific dPCR and quantitative sequencing with dPCR anchoring (relative abundance of a specific taxon measured by sequencing * total 16S rRNA gene copies measured by dPCR). (b) The Log_2_ ratio of the absolute abundance of four bacterial taxa as determined either by taxa-specific dPCR or by quantitative sequencing with dPCR anchoring (N = 32 samples). Data points are overlaid on the box and whisker plot. The body of the box plot represents the first and third quartiles of the distribution and the median line. The whiskers extend from the quartiles to the last data point within 1.5× interquartile range, with outliers beyond. All dPCR measurements are single replicates. (c) Analysis of beta diversity in cecum samples at a series of 10X dilutions (n=1 for each dilution). Mean Aitchison distance for n = 4 sequencing replicates is shown for reference (error bar is standard deviation).

### A ketogenic-diet experiment reveals how a quantitative sequencing framework can provide insights in microbiome studies

To test the impact of using a quantitative framework for 16S rRNA gene amplicon sequencing, we performed a ketogenic-diet study. Our goals were twofold. First, we wished to test whether absolute instead of relative microbial abundances can more accurately quantify changes in microbial taxa between study groups. Second, we wished to investigate how using a quantitative sequencing framework can guide the interpretation of changes in taxa across study conditions. We emphasize that our objective was not to make claims about the effect of a ketogenic diet on the microbiome, but rather to use this model as an illustration of the added benefits of using this quantitative sequencing framework.

After one week on a standard chow diet, 4-week old Swiss Webster mice were split into two groups (n=6 each): one was fed a ketogenic diet and the other a vitamin and mineral matched control diet (Table S3). Stool was sampled immediately before the two diets were introduced (day 0), and again at days 4, 7 and 10. Additionally, on day 10, all mice were euthanized and lumenal and mucosal samples were collected from throughout the GI tract (Fig. 4a). Microbial loads (quantified with dPCR) ranged from ∼10^9^ 16S rRNA gene copies/g in small intestinal mucosa to ∼10^12^ 16S rRNA gene copies/g in stool. On average, we observed lower microbial DNA loads in the mice on the ketogenic diet compared with mice on the control diet, except in the stomach, where loads were similar in mice on both diets (Fig. 4b).

**Figure 4:**
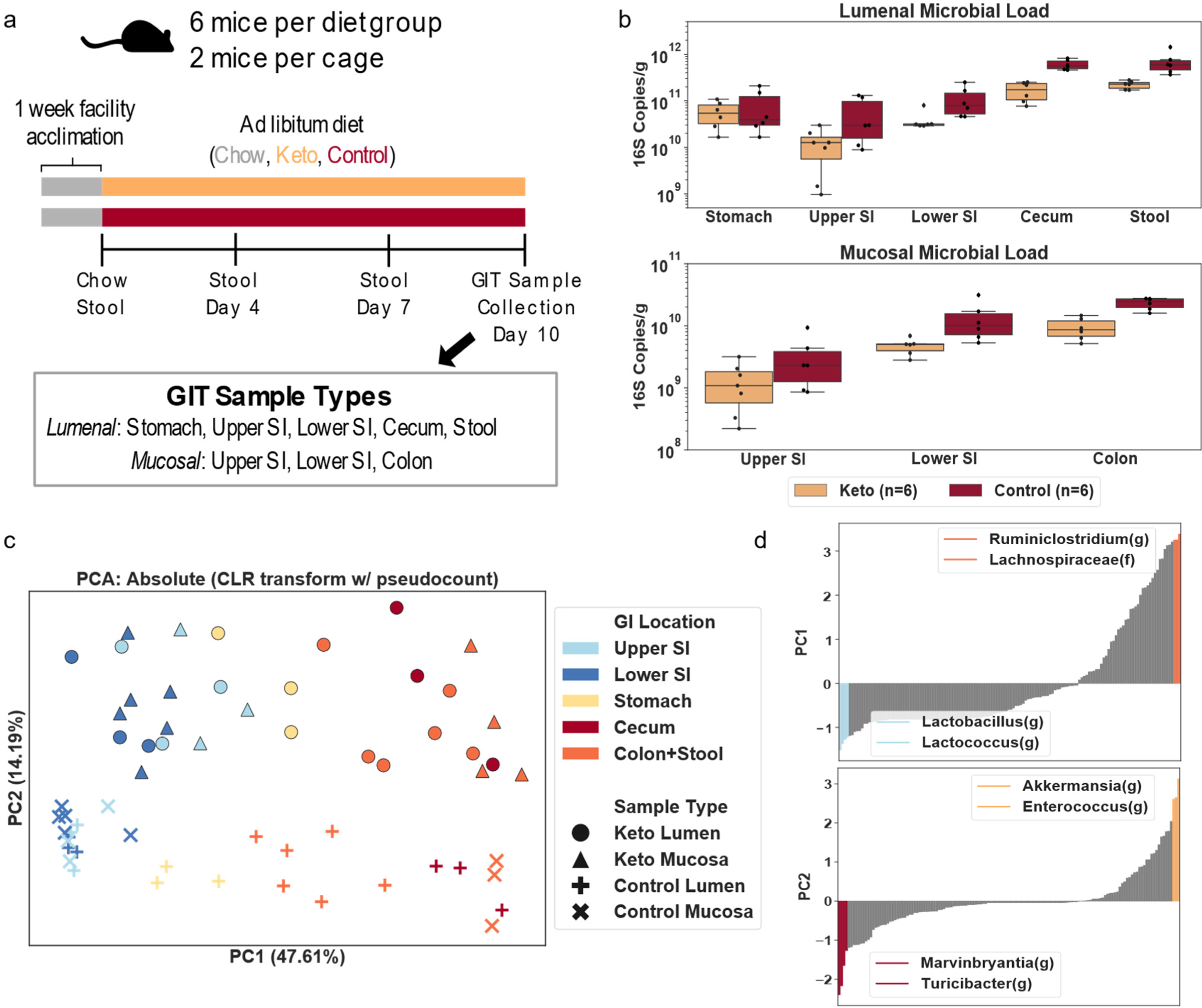
Analysis of data comparing ketogenic and control diets provides changes of total microbial loads, separation of microbial communities by GI location and by diet in principal component analysis, and the top taxa driving the separation of samples along the principal components. (a) Overview of experimental setup and sample-collection protocol. Gastrointestinal tract (GIT) samples were collected from the following regions: stomach, upper small intestine (SI), lower SI, cecum, colon, and stool. (b) Comparison of total microbial loads between ketogenic and control diets in lumenal (top) and mucosal (bottom) samples collected after 10 days on each diet. The body of the box plot represents the first and third quartiles of the distribution and the median line. The whiskers extend from the quartiles to the last data point within 1.5× interquartile range, with outliers beyond. (c) Principal component analysis (PCA) on the centered log-ratio transformed absolute abundances of microbial taxa shows separation by GI location and diet. (d) Ranked order of the eigenvector coefficients scaled by the square root of the corresponding eigenvalue for the top two principal components. The two most positive and most negative taxa are shown.

All stool samples and roughly half of the samples for all other GI sites (evenly distributed across mice on the two diets) underwent 16S rRNA gene amplicon sequencing. Ordination methods (PCA, PCoA, NMDS, etc) are a common exploratory data analysis technique in the microbiome field. Common transformation techniques based on non-Euclidian distances (e.g., Bray-Curtis, UniFrac) can skew the accuracy of visualizations of relative data (Fig S6a).^11^ We used the centered log-ratio transformation (CLR, often used to compute the Aitchison distance) to handle compositional effects, and performed PCA on the transformed absolute abundance data for all samples from the final collection day (Fig. 4c). A clear separation along the first two principal components (PC) was observed. Separation along PC1 was related to the location within the GI tract whereas separation along PC2 was related to the diet. The PCA analysis suggested that stomach samples were distributed somewhere in-between small-intestine and large-intestine samples, possibly resulting from coprophagy in mice.^32, 33^ Additionally, the mucosal and lumenal samples from the small intestine on the control diet seemed to be closer together than on the ketogenic diet (Fig. 4c).

We next investigated which taxa were contributing to separation in our principal component space. We calculated the scaled covariance between each taxon and the first two principal components by multiplying the eigenvectors by the square root of their corresponding eigenvalues. These values are also known as “feature loadings.” Plotting these feature loadings from smallest to highest shows that *Lactobacillus(g)* and *Lactococcus(g)* had the greatest impact on separation along PC1 in the direction of the small intestinal samples whereas *Ruminiclostridium(g)* and *Lachnospiraceae(f)* separated in the direction of the large intestine (Fig. 4d). This matches with what we know about the major genera commonly present in the small and large intestine.^45^ Along PC2 (the “diet axis”), the top two contributing taxa towards the control diet were *Turicibacter(g)* and *Marvinbryantia(g)*, while towards the ketogenic diet *Akkermansia(g)* and *Enterococcus(g)* had the greatest covariance.

Although the CLR transformation preserves distances in principal component space regardless of whether the starting data are relative or absolute, it normalizes out the differences in total loads by looking at log ratios between each taxon’s abundance and the geometric mean of the sample (Fig. S6b). In many cases, we want to know if the absolute load of a taxon is higher or lower under different conditions (e.g., in mice on ketogenic and control diets). When the total microbial load varies among samples, analyses of relative abundance cannot determine which taxa are differentially abundant (Fig. 1). To assess the impact of using absolute quantification in analyses, we analyzed microbiomes of stool samples from mice on ketogenic and control diets. PCA analysis on the CLR-transformed relative abundances of microbial taxa showed separation between the two diets (Fig. 5a). Feature loadings were analyzed as before, but this time total impact of each taxa on the PC space was plotted, which was defined as the sum of the feature loading vectors in PC1 and PC2 (Fig. 5b). The same analysis was performed on the log-transformed absolute abundance data (Fig. 5a). Separation between diets is clear in both relative and absolute abundance analyses, but the contribution of each taxon to the separation differed in direction and magnitude. Comparing the magnitude of feature loadings for two taxa, *Akkermansia(g)* and *Acetatifactor(g)*, between the relative and absolute PCA plots showed obvious differences in the contribution of a given taxa to the separation in principal-component space. Analysis of relative-abundance data implies that *Akkermansia(g)* has the biggest contribution on separation between diets in PC space whereas the absolute abundance data implies that ∼50% of the taxa in the sample have a greater contribution than *Akkermansia(g)* to the separation between the diets in PC space.

**Figure 5:**
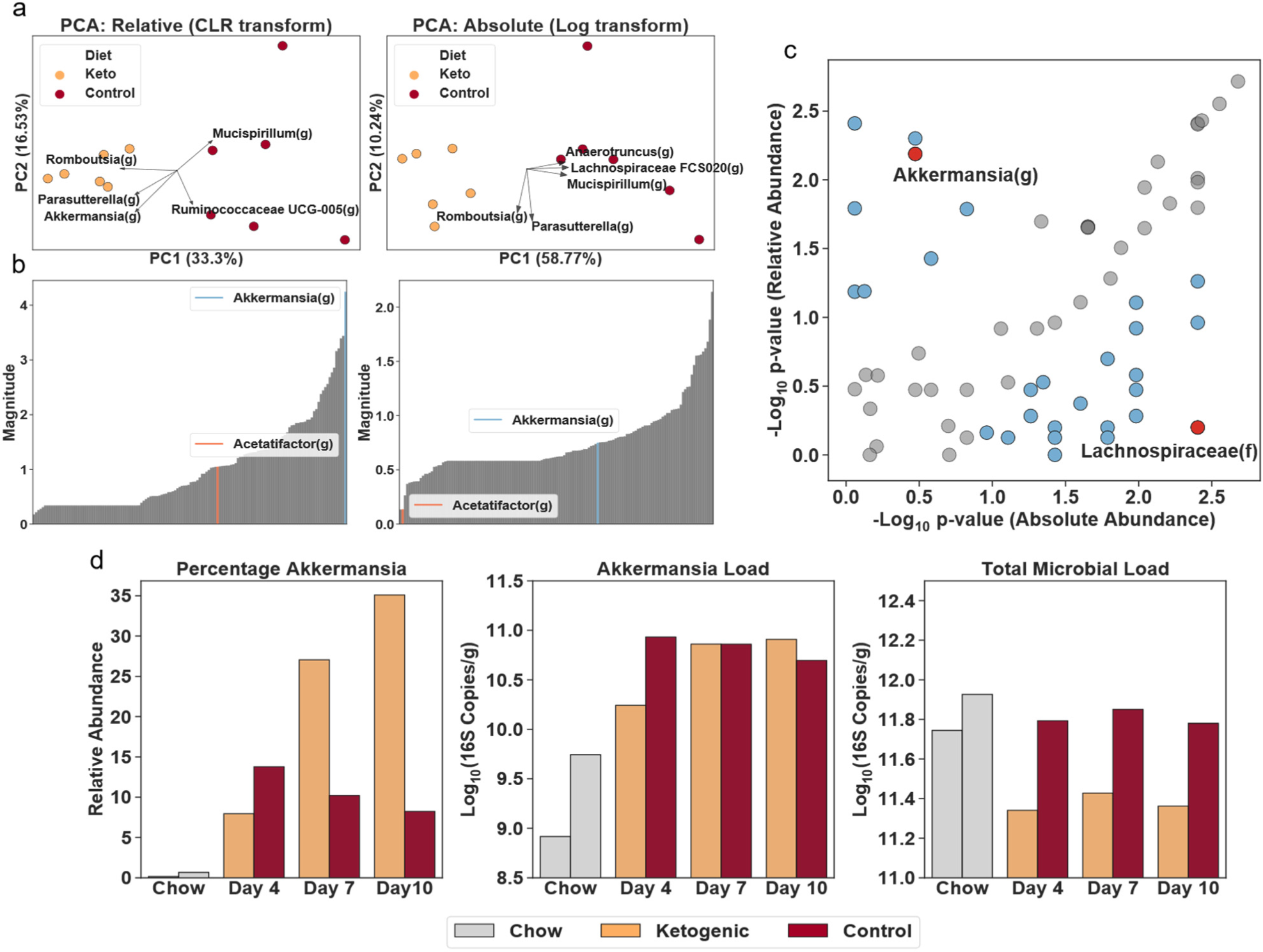
Analyses of relative and absolute microbial abundances from the same dataset result in different conclusions. (a) PCA on centered log-ratio transformed relative abundance data and log transformed absolute-abundance data (only the vectors of the five features with the largest magnitude are shown). (b) The impact of each taxon in the principal-component space (see text for details), with two taxa indicated to illustrate the comparison. (c) A comparison of the taxa determined to be significantly different between diets using relative versus absolute quantification (N = 6 mice per diet). *P-*values were determined by Kruskal-Wallis. Each point represents a single taxon; blue points indicate taxa with the absolute value of *P-*value ratios greater than 2.5; red points indicate two taxa that disagreed significantly between the relative and absolute analyses. (d) For illustrative purposes, a comparison of *Akkermansia(g)* relative abundance (percentage of Akkermansia), absolute abundance (Akkermansia load), and total microbial load between stool samples from one mouse on each diet.

PCA is only an exploratory data-analysis technique, so we next used a non-parametric statistical test to test for differentially abundant taxa in stool samples from mice on control and ketogenic diets (Fig. 5c).^46^ We performed separate analyses of the relative and absolute abundance data. We plotted the −log10 *P*-value for each taxon’s relative abundances against the corresponding −log10 *P*-value for that taxon’s absolute abundances. Points along the diagonal indicate congruence between the predictions from the relative and absolute abundance data. Points in the upper left corner indicate taxa that differed between the diets in the analysis of relative-abundance but not in the analysis of absolute abundance. Conversely, points in the lower right corner indicate taxa that do not differ between diets in the analysis of relative abundance but do differ in the analysis of absolute abundance. *Akkermansia(g)* is an example of a microbe that appears to differ (*P* = 6.49 × 10^-3^) between mice on the two diets in the relative-abundance analysis but not in the absolute-abundance analysis (*P* = 3.37 × 10^-1^). *Lachnospiraceae(f)* showed the opposite trend; in the relative-abundance analysis it appears unchanged (*P* = 6.31 × 10^-1^) but in the absolute-abundance analysis it differs (*P* = 3.95 × 10^-3^) between the two diets. Neither of these analyses is wrong, they are simply asking two different questions: with relative data, the question is whether the percentage of that microbe is different between two conditions whereas with absolute data, the question is whether the abundance of that microbe is different between two conditions.

To explore one example of how different interpretations of how taxa differ between study conditions occur when using relative versus absolute abundance, we analyzed *Akkermansia(g)* in stool across each of the three time points on experimental diets (days 4, 7, and 10) and day 0 on chow diet. For simplicity in this illustration, we compared data from one mouse on each diet, but the trends hold on average between all mice on the two diets (Fig. S7). Analysis of relative microbial abundance demonstrated ∼3X higher abundance of *Akkermansia(g)* in samples from the ketogenic compared with the control diet on days 7 and 10. However, when analyzing the difference in absolute abundance, more nuanced conclusions emerged. The rise in *Akkermansia(g)* results from switching mice from chow to experimental diets. The resulting *Akkermansia(g)* loads are similar in the two diets on days 7 and 10. However, the ketogenic diet reduces the total microbial load relative to both chow and control diets, therefore leading to the observed higher % of *Akkermansia(g)* in samples from mice on ketogenic diet.

### Absolute quantification accurately reveals the direction and magnitude of changes in microbial taxa

We next analyzed the absolute microbiota abundances in stool and lower small intestinal mucosa samples from day 10. A volcano plot, akin to those used in gene expression studies, was used to represent the overall changes in taxa abundances between the two diets, and the absolute abundance of each taxon was indicated by the size of its symbol (Fig. 6a). *P*-values from the Kruskal-Wallis tests were corrected for multiple hypothesis testing with the Benjamini– Hochberg method, resulting in q-values.^46, 47^ A false discovery rate (FDR) of 10% was labeled on the volcano plot and q-values < 0.1 were used as a cutoff for designating differential taxa for downstream analyses. Comparisons between the two GI locations showed substantial differences in microbial response to diet by location. In stool, approximately 66% of the differential taxa were lower on the ketogenic diet vs the control diet whereas in the lower SI mucosa, > 80% of the differential taxa were more abundant in the ketogenic diet than control diet (Table S4, Table S5).

**Figure 6:**
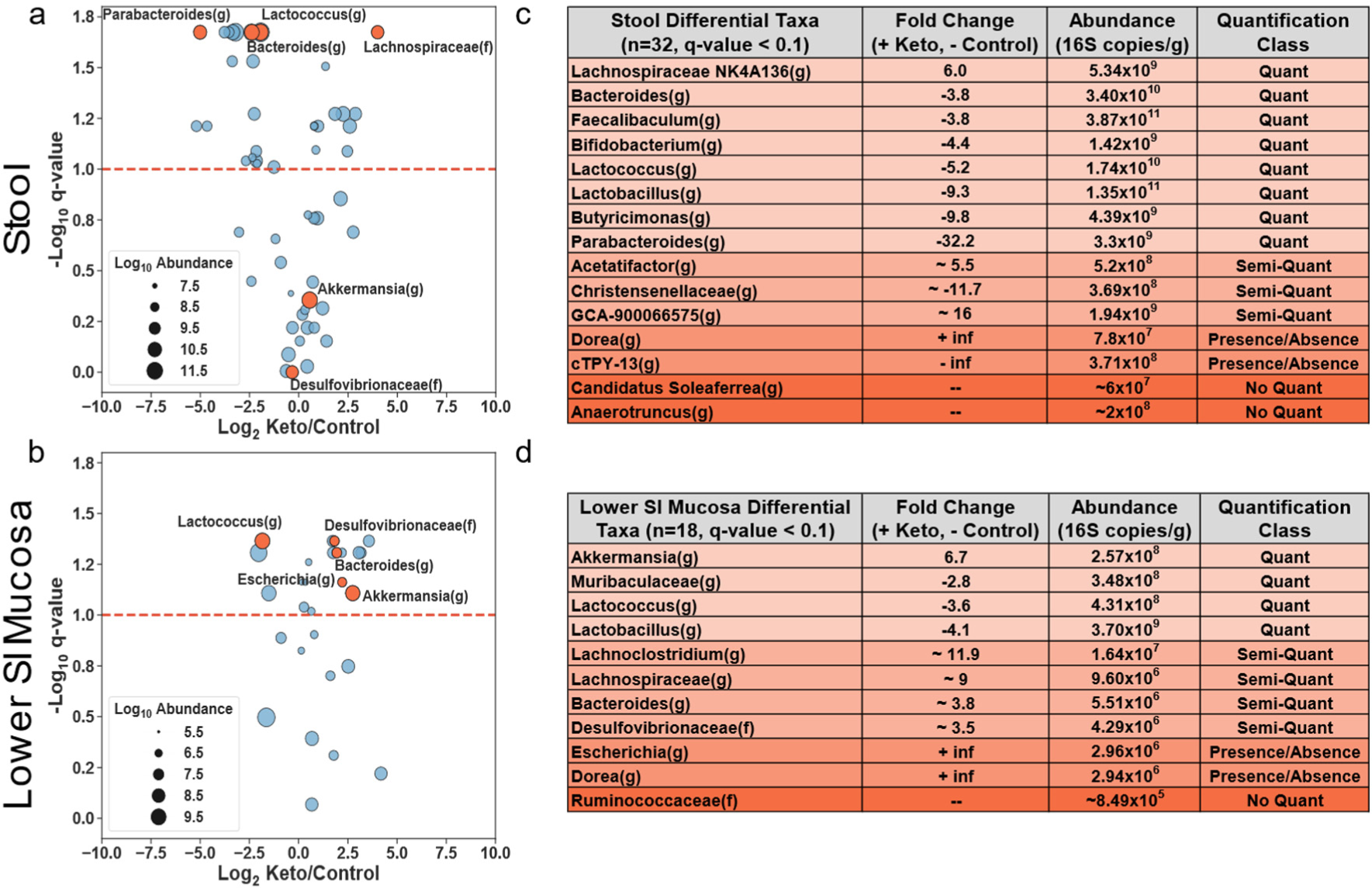
A quantitative framework that explicitly incorporates limits of quantification separates differential microbial taxa into four classes, and for each GI location identifies a distinct set of differential taxa, including taxa with opposite patterns in stool and SI mucosa. (a-b) Microbial taxa in stool (a) or lower small-intestine (b) mucosa in mice on ketogenic (N = 6) and control (N = 6) diets. The fold change on the x-axis is the Log_2_ ratio of the average absolute loads of taxon loads in each diet. Negative values indicate lower loads in ketogenic diet compared to control diet. The q-value for a taxon indicates the significance of the difference in absolute abundances between the two diets and were obtained by Kruskal-Wallis with a Benjamini–Hochberg correction for multiple hypothesis testing. The Log_10_ absolute abundance of each taxon is indicated by circle size. Orange circles indicate taxa discussed in the main text including taxa that show discordant fold changes between stool and lower SI mucosa. The red dashed line is shown at a q-value representing a 10% false-discovery rate. (c-d) A subset of taxa from stool (c) and lower SI mucosa (d) that were significantly different between diets (q-values < 0.1) and their corresponding fold change, absolute abundance (larger of the average absolute abundances between the two diets), and quantification class. Quantification class is determined by whether one or both measurements were above or below the lower limit of quantification and the limit of detection.

Next, we highlighted several specific differential taxa that were discordant between stool and lower SI mucosa. (1) *Bacteroides(g)* was lower on ketogenic diet in stool and higher on ketogenic diet in lower SI mucosa. This type of result could lead researchers who analyze stool samples to believe that lower levels of *Bacteroides(g)* may be associated with a phenotype when it could be the opposite if the phenotype is driven by the SI mucosal microbiota. (2) *Parabacteroides(g)* and *Lachnospiraceae GCA-900066575(g)* showed the highest fold changes (in opposite directions) in stool but were not detected in the lower SI mucosa. The opposite was observed for *Escherichia(g)*, which was more abundant in the ketogenic diet than the control diet in the lower SI mucosa but was not detected in stool. (3) *Akkermansia(g)* and *Desulfovibrionaceae(f)* were more abundant in the ketogenic diet than the control diet in the lower SI mucosa but were similar between the two diets in stool. Such microbes could have a relationship with phenotype through the small intestine but would be missed if only stool samples are analyzed.

A further breakdown of the differential taxa, using our quantitative limits of sequencing accuracy (defined earlier), allowed us to categorize four distinct scenarios that describe how microbes differed between GI locations of mice on the two diets. We refer to these four scenarios as “quantification classes” (Fig. 6b). First, there were microbes that were present in one diet and absent in the other (“presence/absence” class). For example, *Dorea(g)*, in stool, and *Escherichia(g)*, in SI mucosa, were absent from the control diet but present in the ketogenic diet. Second, there were microbes above the detection limit but below the quantitative limit in both diets (“no quant” class). For example, in stool, *Candidatus Soleaferrea(g)*, ranges in relative abundance from 0.002% to 0.025%, well below the 30% CV quantification threshold of 0.04% (as defined in Fig. 2d). Thus, we cannot quantitatively define the difference of this microbe between mice on the two diets. Third, microbes were above the detection limit in both diets but only above the quantitative limit in one of the diets (“semi-quant” class). For example, *Desulfovibrionaceae(f)* in the lower small-intestine mucosa was above the detection limit in mice on both diets but only above the quantitative limit in mice on the ketogenic-diet, so although we can be confident that a difference between the diets exists, we cannot be confident in our measurement of the magnitude of that difference. Fourth, microbes were found above the quantitative limits in both diets (“quant” class). For example, for *Parabacteroides(g)* in stool, we can be confident in both the difference between the diets (it was more abundant in the control diet) and in the magnitude of that difference (a 32.2-fold difference). We have the lowest confidence in the measured absolute fold change of a taxon that is classified in the presence/absence class, and the greatest confidence in a taxon in the quant class.

## Discussion

In this study, we have shown that this technology performs across biogeographically diverse samples with microbial loads spanning over 6 orders of magnitude. Our lower limits of quantification for total microbial load from lumenal (e.g., stool, cecum contents) and mucosal samples were 4.2 × 10^5^ 16S rRNA gene copies/g and 1.0 × 10^7^ 16S rRNA gene copies/g respectively. These lower limits were mainly restricted by the column-based extractions used which require < 200 mg of sample input for lumenal contents and < 8 mg of input for mucosal samples. This sample input is limited by the high concentration of PCR inhibitors and host DNA in these samples. New sample-processing methods that deplete host DNA before extraction (e.g. the use of propidium monoazide (PMA) or saponin with DNase)^48, 49^ could help improve the quantitative limits in samples with high levels of host DNA (e.g., mucosa) by removing non-microbial DNA before extraction. Such host-depletion methods could also improve performance of other current or future methods of quantitative sequencing. Before these methods are introduced into quantitative sequencing protocols, they will require extensive validation to understand the impacts host DNA depletion has on the microbial load and composition of these samples, which will affect the accuracy of any absolute-abundance technique. We showed that the precision of any individual taxon’s abundance can be defined as a function of that taxon’s relative abundance and the sequencing depth. These accuracy thresholds generally state that all taxa with relative abundance > 0.01% have a maximum %CV of 30%. We did not quite reach the theoretical limit of Poisson precision (Fig. 2c), which might be explained by slight differences in PCR amplification between high- and low-abundance microbes, and could potentially be corrected with single-molecule counting techniques utilizing unique molecular identifiers (UMIs).^50, 51^ Interestingly, the precision of these abundance measurements did not differ between high input DNA samples (1.2 × 10^7^ 16S rRNA gene copies) and low-input DNA samples (1.2 × 10^4^ 16S rRNA gene copies), even though the low-input sample required 10 additional PCR cycles. The lack of an increase in observed chimeric sequences in the low-input sample indicates that PCR bias from chimera generation may occur mainly during over-amplification; thus, we suggest monitoring library-prep amplification reactions with qPCR and stopping reactions during the late exponential phase.

Our quantitative sequencing method, as validated, is subject to some of the same limitations of general 16S rRNA gene amplicon sequencing. Primarily, the accuracy of any given taxon’s abundance is believed to be impacted by amplification bias. We showed that the abundances of *Akkermansia muciniphila(s), Lachnospiraceae(f), Bacteroidales(o)*, and *Lactobacillaceae(f)* could be quantified with similar precision (2X), but different accuracy, i.e. *Akkermansia muciniphila(s)* abundance was ∼2.5X higher in the quantitative-sequencing estimate compared with the estimate from dPCR with taxa-specific primers. This offset was consistent between samples, indicating that it may be related to differences in primer coverage between the taxon-specific primer set and the universal primer set used in this study. Nevertheless, such offsets should be similar if the same library-prep conditions are used, so one can reliably compare taxa among groups or studies and the use of UMIs may further eliminate any potential amplification biases. We note that dPCR-based total microbial load measurements should be more robust to amplification biases of individual taxa. Additionally, the total microbial load measurement will be affected by the 16S rRNA primer set chosen and its respective taxonomic coverage. The primers in this study were chosen to have broad coverage and also to limit amplification of host mitochondrial DNA,^31-33^ to ensure proper quantification of mucosal and small-intestine samples with high host DNA loads. Finally, to take full advantage of the power of this quantitative framework, study designs must incorporate proper sampling techniques to address spatiotemporal variation in microbial abundances.^22^

A method-specific limitation is the requirement of an additional step, dPCR total microbial load quantification, which consumes a portion of the extracted DNA sample. This limitation is minor because dPCR generally requires at least 100 copies for a measurement with a ∼10% Poisson error, which is much less than the roughly 10,000 copies required for sequencing. Additionally, the absolute abundances are reported in 16S rRNA gene copies/g and require conversion to number of cells/g, which has standard limitations (e.g., the completeness of rRNA databases and copy-number variation among similar species). However, when comparing taxa across study groups, the 16S rRNA gene copies per taxonomic group should be similar. Finally, this method was only validated for 16S rRNA gene amplicon sequencing; thus, further validation would be required for applying this method to converting metagenomic sequencing from relative to absolute quantification.

We applied the quantitative framework to a murine ketogenic-diet study to identify how microbial taxa at several GI locations respond to diet. Because total microbial loads were lower in the ketogenic diet compared to the control, analysis of absolute abundance was required to correctly identify differential taxa. The lower load observed on the ketogenic diet can likely be explained by its lower fiber and carbohydrate content, as these dietary components are main substrates for many gut microbes.^52^ Many factors (including diet) that induce changes in relative microbial abundances can also impact total microbial load.^25, 53^ Even among healthy mice on the same (chow) diet, total microbial loads in stool can differ by 10 times.^25^ Such variation in total microbial load likely contributes to the noise in microbiome studies. Another insight of this study was that we found different patterns in the microbial communities at each GI sampling site. For example, *Akkermansia(g)* loads did not differ between diets in stool, but they were significantly greater in the small-intestine mucosa in the ketogenic diet compared with the control. *Bacteroides(g)* load was lower in stool and greater in the small-intestine mucosa in the ketogenic relative to the control diet. Clearly, differential taxa at one GI location cannot be used as a proxy for measuring differential taxa at another GI location. To our knowledge, this is the first microbiome study to show that microbial taxa in the small intestine and the stool can change in different directions and by different magnitudes in response to diet. Furthermore, for each taxon, this method enables a comparison of absolute microbial abundance to limits of detection and quantification. This comparison separates differential taxa into four classes (Quant, Semi-Quant, No Quant, Presence/Absence) which provide a convenient shortcut for more quantitative interpretation of microbiome studies. It should be noted that the absence of a microbe in a dataset is a factor of the sequencing depth, and just because a microbe is not found in the sequencing data does not mean it is not in the sample. However, with absolute anchoring, one can confidently say that when a microbe is not found, that microbe is below a given abundance.

We have not focused on correlations among taxa in this dataset. However, the absolute abundance measurements acquired using our method should help overcome many of the limitations of correlation-based analyses on relative abundances^54, 55^ and enable analyses using standard methodologies like Spearman’s rank correlation (Fig. S8). However, further work will be required to properly address the impact that correlations between total microbial loads will have on taxon-based correlation networks. In addition, new statistical and/or experimental design methods may be required for interpreting the correlations between a taxon’s presence and/or total load and observed phenotypes.

This method overcomes three bottlenecks to wider adoption of absolute quantitative measurements in microbiome analysis: (i) performance across samples with a wide range of microbial loads; (ii) performance across biogeographically diverse sample types (iii) explicit evaluation of limits of quantification of the method. This method will be useful in other areas that benefit from quantitative analysis, such as monitoring microbial communities during manufacturing of complex probiotic mixtures^56^ and monitoring changes of host-associated microbial communities over time (e.g. in health, aging and development, disease progression, and during probiotic or other treatments). Applying absolute quantification^19-21, 23-28, 32, 33^ of microbial taxa to biogeographically relevant GI locations will provide researchers with new insights in how microbial communities affect host phenotypes.

## Methods

### Mice

All animal husbandry and experiments were approved by the Caltech Institutional Animal Care and Use Committee (IACUC protocols #1646 and #1769). Male and female germ free (GF) C57BL/6J mice were bred in the Animal Research Facility at Caltech, and 4-week-old female specific-pathogen-free (SPF) Swiss Webster mice were obtained from Taconic Farms (Germantown, NY, USA). Experimental animals were fed standard chow (Lab Diet 5010), 6:1 ketogenic diet (Harlan Teklad TD.07797; Table S3) or vitamin- and mineral-matched control diet (Harlan Teklad TD.150300; Table S3). Diet design and experimental setup were taken from a recently published study.^40^ To minimize cage effects, mice were housed two per cage with three cages per diet group. Custom feeders, tin containers approximately 2.5 inches tall with a 1-inch diameter hole in the top, were used for the ketogenic diet as it is a paste at room temperature. Mice were euthanized via CO_2_ inhalation as approved by the Caltech IACUC in accordance with the American Veterinary Medical Association Guidelines on Euthanasia.^57^

### Microbial Samples

The mock microbial community (Zymobiomics Microbial Community Standard; D6300) was obtained from Zymo Research (Irvine, CA, USA). This community is stored in DNA/RNA Shield, which interferes with extraction efficiency at high concentrations (data not shown). We found that a 100 µL input of a 10X dilution of the microbial community stock is the maximum input that the Qiagen DNeasy Powersoil Pro Kit can handle without recovery losses. Negative control blanks were also used which included 100 µL of nuclease free water instead of mock community.

Fresh stool samples were collected immediately after defecation from individual mice and all collection occurred at approximately the same time of day. For intestinal samples, the GIT was excised from the stomach to the anus. Contents from each region of the intestine (stomach, upper half of SI, lower half of SI, cecum, and colon) were collected by longitudinally opening each segment with a scalpel and removing the content with forceps. Terminal colonic pellets are referred to as stool. After contents were removed the intestinal tissue was washed by vigorously shaking in cold sterile saline. The washed tissue was placed in a sterile petri dish and then dabbed dry with a Kimwipe (VWR, Brisbane, CA, USA) before scraping the surface of the tissue with a sterile glass slide. These scrapings were collected as the mucosa samples. All samples were stored at −80 °C after cleaning and before extraction of DNA.

### DNA Extraction

DNA was extracted from all samples by following the Qiagen DNeasy Powersoil Pro Kit protocol (Qiagen; Valencia, CA, USA). Bead-beating was performed with a Mini-BeadBeater (BioSpec, Bartlesville, OK, USA) for 4 min. To ensure extraction columns were not overloaded, we used ∼10 mg of scrapings and ∼50 mg of contents. Half of the lysed volume was loaded onto the column and elution volume was 100 µL. Nanodrop (NanoDrop 2000, ThermoFisher Scientific) measurements were performed with 2 µL of extracted DNA to ensure concentrations were not close to the extraction column maximum binding capacity (20 µg).

### Absolute Abundance

The concentration of total 16S rRNA gene copies per sample was measured using the Bio-Rad QX200 droplet dPCR system (Bio-Rad Laboratories, Hercules, CA, USA). The concentration of the components in the dPCR mix used in this study were as follows: 1x EvaGreen Droplet Generation Mix (Bio-Rad), 500 nM forward primer, and 500 nM reverse primer. Universal primers to calculate the total 16S rRNA gene concentrations were a modification to the standard 515F-806R primers^4^ to reduce host mitochondrial rRNA gene amplification in mucosal and small-intestine samples (Table S6).^31-33^ Thermocycling for universal primers was performed as follows: 95 °C for 5 min, 40 cycles of 95 °C for 30 s, 52 °C for 30 s, and 68 °C for 30 s, with a dye stabilization step of 4 °C for 5 min and 90 °C for 5 min. All ramp rates were 2 °C per second. The concentration of taxon-specific gene copies per sample was measured using a similar dPCR protocol, except with different annealing temperatures. Annealing temperatures during thermocycling for taxa-specific primers can be found in Table S6. The concentration of total 16S rRNA gene copies per sample was also estimated using qPCR with the CFX96 RT-PCR machine (Bio-Rad). The concentration of the components in the qPCR mix used in this study were as follows: 1x SsoFast EvaGreen Supermix (BioRad), 500 nM forward primer, and 500 nM reverse primer. Thermocycling was performed as follows: 95°C for 3 min, 40 cycles of 95 °C for 15 s, 52 °C for 30 s, and 68 °C for 30 s. All dPCR measurements are single replicates.

Concentrations of 16S rRNA gene per microliter of extraction were corrected for elution volume and losses during extraction before normalizing to the input sample mass (Equation 1).

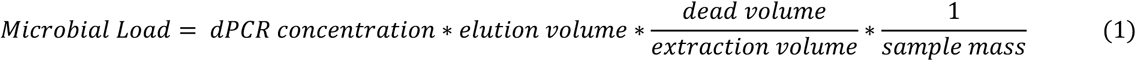

Absolute abundance of individual taxa was calculated either by dPCR with taxa-specific primers or multiplying the total microbial load from Eqn. 1 by the relative abundance from 16S rRNA gene amplicon sequencing.

### 16S rRNA Gene Amplicon Sequencing

Extracted DNA was amplified and sequenced using barcoded universal primers and protocol modified to reduce amplification of host DNA^31-33^. The variable 4 (V4) region of the 16S rRNA gene was amplified in triplicate with the following PCR reaction components: 1X 5Prime Hotstart mastermix, 1X Evagreen, 500 nM forward and reverse primers. Input template concentration varied. Amplification was monitored in a CFX96 RT-PCR machine (Bio-Rad) and samples were removed once fluorescence measurements reached ∼10,000 RFU (late exponential phase). Cycling conditions were as follows: 94 °C for 3 min, up to 40 cycles of 94 °C for 45 s, 54 °C for 60 s, and 72 °C for 90 s. Triplicate reactions that amplified were pooled together and quantified with Kapa library quantification kit (Kapa Biosystems, KK4824, Wilmington, MA, USA) before equimolar sample mixing. Libraries were concentrated and cleaned using AMPureXP beads (Beckman Coulter, Brea, CA, USA). The final library was quantified using a High Sensitivity D1000 Tapestation Chip. Sequencing was performed by Fulgent Genetics (Temple City, CA, USA) using the Illumina MiSeq platform and 2×300bp reagent kit for paired-end sequencing.

### Data Analysis and Statistics

#### 16S rRNA Gene Amplicon Data Processing

Processing of all sequencing data was performed using QIIME 2 2019.1.^58^ Raw sequence data were demultiplexed and quality filtered using the q2-demux plugin followed by denoising with DADA2.^59^ Chimeric read count estimates were estimated using DADA2. Beta-diversity metrics (Aitchison distance,^9^ Bray-Curtis Dissimilarity) were estimated using the q2-diversity plugin after samples were rarefied to the maximum number of sequences in each of the relevant samples. Rarefaction was used to force zeros in the dataset to have the same probability (across samples) of arising from the taxon being at an abundance below the limit of detection. Although rarefaction may lower the statistical power of a dataset^60^ it helps decrease biases caused by different sequencing depths across samples.^12^ Taxonomy was assigned to amplicon sequence variants (ASVs) using the q2-feature-classifier^61^ *classify-sklearn* naïve Bayes taxonomy classifier against the Silva^62^ 132 99% OTUs references from the 515F/806R region. All datasets were collapsed to the genus level before downstream analyses. All downstream analyses were performed in IPython primarily through use of the *Pandas, Numpy* and *Scikit-learn* libraries.

#### Data Transforms and Dimensionality Reduction

For dimensionality reduction techniques requiring a log transform, a pseudo-count of 1 read was added to all taxa. With relative abundance data, the centered log-ratio transform was used (Equation 2) to handle compositional effects whereas a log transform was applied to the absolute-abundance data to handle heteroscedasticity in the data.

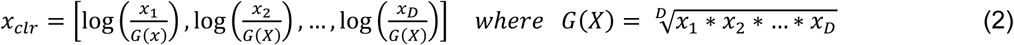

For comparative purposes, principal co-ordinates analysis (PCoA) was also performed using the Bray-Curtis dissimilarity metric. Principal component analysis (PCA) and PCoA were performed using *scikit-learn* decomposition methods. Feature loadings for each principal component were calculated by multiplying each eigenvector by the square root of its corresponding eigenvalue. All data were visualized using matplotlib and seaborn.

#### Taxa Limits of Quantification

Poisson confidence intervals were calculated by bootstrapping Poisson samples for rate parameters across the percentage abundance range (0–1) corresponding to either the input DNA copies or number of reads. We took 10^4^ bootstrap replicates with a Poisson sample size of 4 to match the number of replicates we sequenced. The %CV for each replicate was calculated and the middle 95^th^ percentile was shown as the confidence interval.

Thresholds for percentage abundance were calculated by first fitting a negative exponential curve *y = ax*^−*b*^ to the plot of %CV versus percentage abundance using SciPy. Then the percentage abundance at a given %CV threshold was determined. This process was repeated after subsampling the data at decreasing read depths to find the relationship between percent abundance accuracy limits at sequencing depth.

#### Measurement Uncertainty

When measuring the absolute abundance of a given taxon in a sample, many factors contribute to the uncertainty of the measurement. Two primary factors, extraction efficiency and average amplification efficiency for each taxon, should be equivalent for each taxon across samples processed under identical conditions and thus neither should impact the discovery of differential taxa. However, other factors contributing to the uncertainty of an absolute-abundance measurement vary among samples and can impact the discovery of differential taxa. At least six independent errors can contribute to the overall uncertainty of a taxon’s absolute abundance: (i) extraction error (ii) the Poisson sampling error of dPCR, (iii) the Poisson sampling error of sample input into an amplification reaction to make a sequencing library, (iv) the uncertainty in the amplification rates among sequences, (v) the Poisson sampling error of the sequencing machine, and (vi) the uncertainty in taxonomic assignment resulting from different software programs that differ in how they convert raw sequencing reads to a table of read counts per taxon.

To measure the total error in our absolute-abundance measurements, we compared the true absolute load value of four “representative” taxa (taxa that are common gut flora from different taxonomic ranks) as measured by taxa-specific dPCR, with the value obtained from our method of quantitative sequencing with dPCR anchoring (Fig. 3b) and then analyzed the relative error in these measurements, defined as the log_2_ of the observed taxon load over the true taxon load. We constructed a quantile-quantile (Q–Q) plot (Fig. S9) of the mean-centered log_2_ relative errors and found that the errors appeared normally distributed. We confirmed this by running a Shapiro–Wilk test (*P*-value = 0.272) on the mean-centered log_2_ relative errors, which uses a null hypothesis that the dataset comes from a normal distribution. The standard deviation of the mean-centered log_2_ relative errors was 0.48, which results in a 95% confidence interval of ∼(−1,1), indicating a 2x precision on each individual measurement. However, as seen with *Akkermansia(g)* (Fig. 3b), accuracy offsets may exist for specific taxa. It is important to note that all samples used in this analysis had relative abundances above the 50% CV threshold defined in Fig. 2d and thus we do not make any conclusions about the precision of absolute abundance measurements for taxa with relative abundances below the 50% CV threshold.

#### Biological Uncertainty and Statistical Inference Methods

When measuring the absolute abundance of a taxon from a defined population (e.g., healthy adults, mice on a ketogenic diet) it is unlikely this abundance comes from a well-defined statistical distribution. Given this inherent limitation, we used non-parametric statistical tests, which do not rely on distributional assumptions, for our differential abundance analyses.

Statistical comparisons between diet groups were analyzed using the Kruskal–Wallis^46^ rank sums test with Benjamini– Hochberg^47^ multiple hypothesis testing correction. All statistical tests were implemented using *SciPy*.*stats Kruskal* function and *statsmodels*.*stats*.*multitest multipletests* function with the *fdr_bh* option for Benjamini-Hochberg multiple-testing correction. When calculating differentially abundant taxa, only taxa present in at least 4 out of 6 mice in a group were considered to remove fold-change outliers when plotting (Fig. 6a-b).

#### Correlation Analysis

Samples were separated by diet (ketogenic and control) and only stool samples were used (days 4, 7, and 10). The total microbial load and top 30 taxa with the highest average absolute abundance were selected for analysis. Spearman’s rank correlation coefficient and corresponding *P*-values were calculated for all pairwise interactions using the *scipy*.*stats*.*spearmanr* function. Benjamini–Hochberg procedure was to calculate q-values, which account for multiple hypothesis testing. A heatmap of the diagonal correlation matrix was plotted (Fig. S8) for q-values <10% FDR.

## Data Availability

The complete sequencing data generated during this study are available in the National Center for Biotechnology Information Sequence Read Archive repository under study accession number PRJNA575097. Raw data for all figures available through CaltechDATA: https://data.caltech.edu/records/1371.

## Acknowledgements

This work was supported in part by the Kenneth Rainin Foundation (2018-1207), the Army Research Office (ARO) Multidisciplinary University Research Initiative (MURI #W911NF-17-1-0402), and a National Institutes of Health Biotechnology Leadership Pre-doctoral Training Program (BLP) fellowship from Caltech’s Donna and Benjamin M. Rosen Bioengineering Center (T32GM112592, to J.T.B.). We thank Elaine Hsiao and Christine Olson for helpful discussions and input on the experimental design and diets; we thank the Caltech Bioinformatics Resource Center for assistance with statistical analyses; we acknowledge the Caltech animal facility for experimental resources; we thank the Caltech Office of Laboratory Animal Resources and the veterinary technicians at Caltech for technical support; and Natasha Shelby for contributions to writing and editing this manuscript.

## Author Contributions

JTB: validated limits of digital PCR assay with mock microbial communities in germ-free tissues; designed, performed, and analyzed experiments to validate accuracy of quantitative sequencing with dPCR anchoring; established the quantitative limits of an individual taxon’s absolute abundance; conducted the ketogenic animal study; analyzed all data; created all figures; and wrote the paper.

SRB: co-developed the idea of quantitative sequencing with dPCR anchoring for absolute quantification of total microbial loads and taxa absolute abundances in lumenal and mucosal samples; contributed the method for quantitative sequencing with dPCR anchoring in lumenal and mucosal samples; contributed ideas and provided support for animal study design; contributed ideas for data analysis and representation.

RFI: contributed to study design and manuscript preparation.

