## Supplemental Information for "A Quantitative Sequencing Framework for Absolute Abundance Measurements of Mucosal and Lumenal Microbial Communities"

Figures S1-S9

Tables S1-S6

Supplementary References

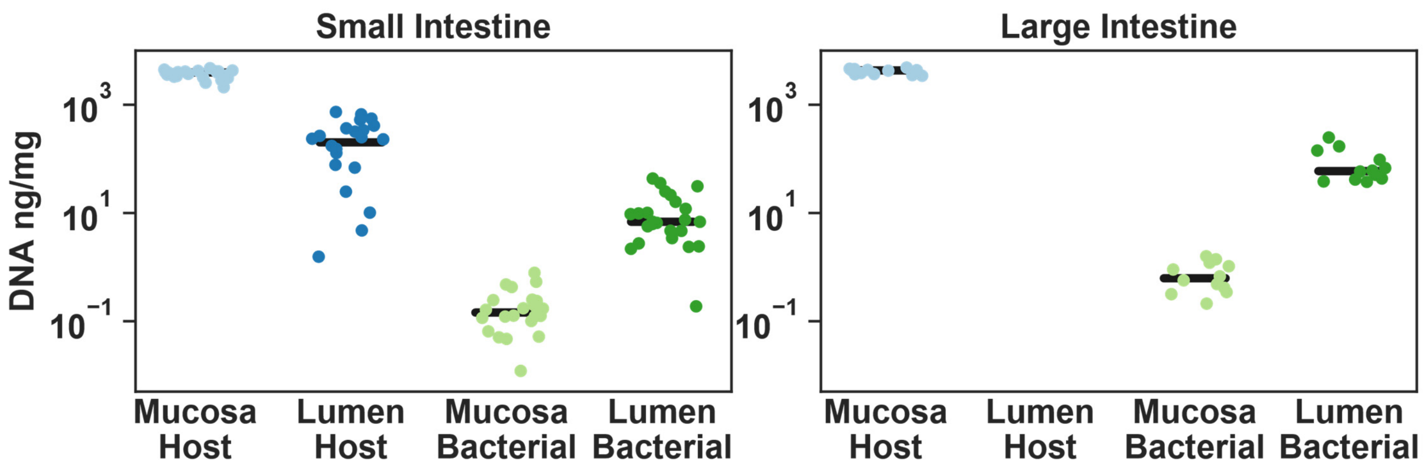


**Figure S1: Total DNA loads in small intestine and large intestine mucosa and lumen.** Extracted DNA samples from mice in the ketogenic-diet group were measured by Nanodrop (total DNA) and digital PCR (microbial DNA). The horizontal lines represents the means and the points represent individual biological replicates (N = 24 for small intestine; N = 12 for large intestine).

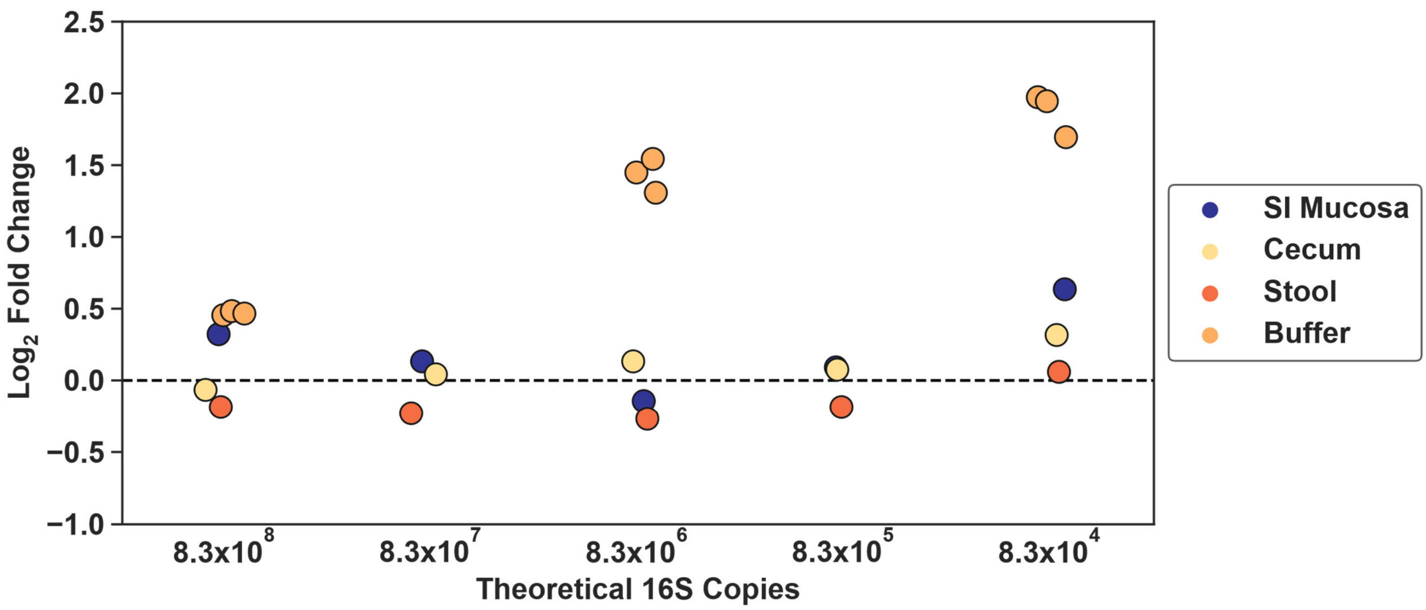


**Figure S2: Extraction and total DNA measurement accuracy of an eight-member mock microbial community dilutions spiked into extraction buffer or small-intestine mucosa, cecum, or stool from germ free mice.** $Log_2$ fold change between theoretical and dPCR measured copies of 16S rRNA gene after extraction with varying input levels. Three technical replicates for buffer extractions are shown. All other sample types shown are N = 1 to illustrate the biological noise among sample types.

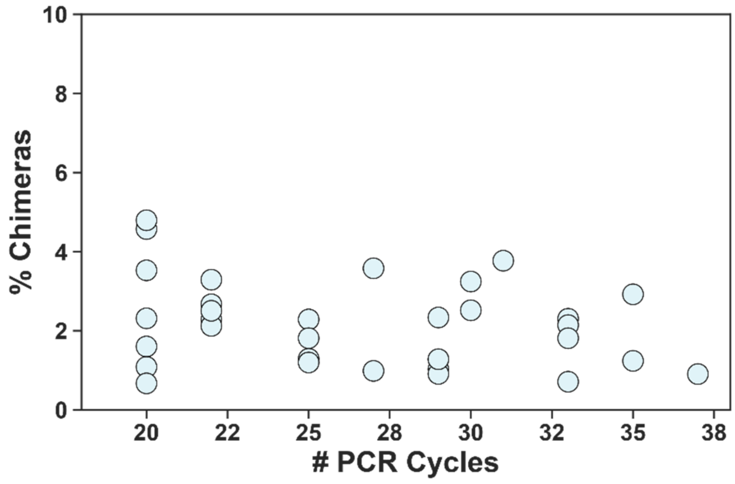


**Figure S3: Chimeric sequence prevalence is not determined by the number of PCR cycles.** Relationship between the number of PCR cycles during the amplification reaction for library prep and the percentage of chimeric sequences detected by Divisive Amplicon Denoising Algorithm 2 (DADA2).[1] N = 33 samples that were sequenced from mice in the ketogenic-diet group.

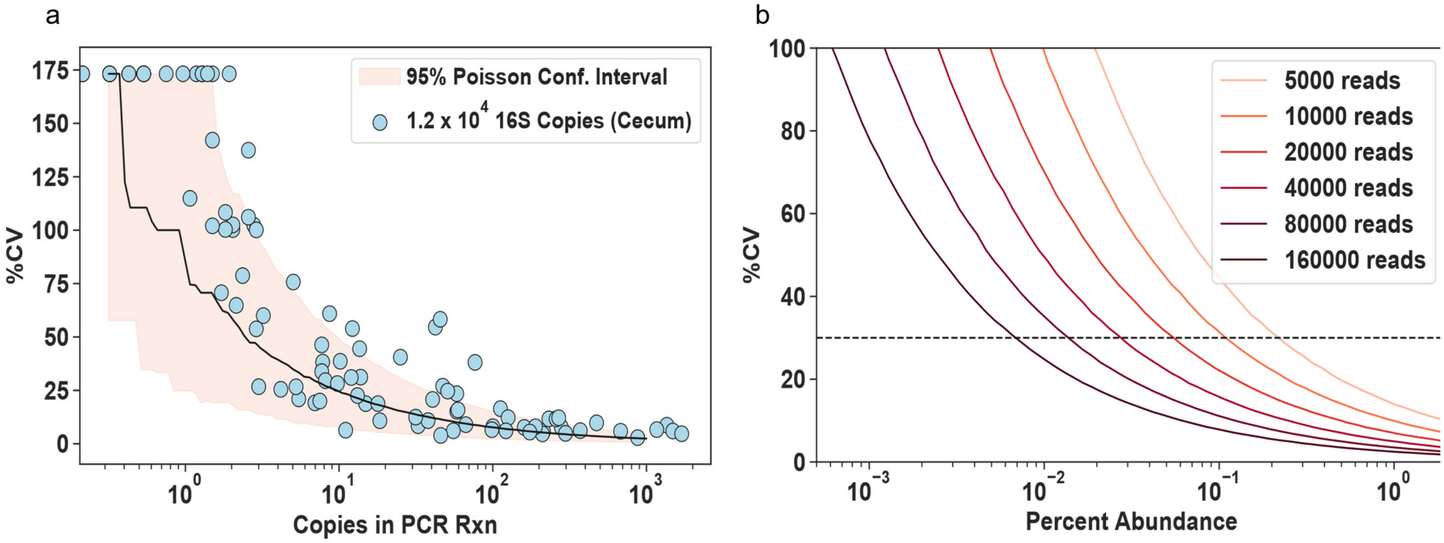


**Figure S4: Poisson limits of sequencing accuracy.** (a) Relationship between the relative abundance of each taxon and % coefficient of variation (CV) using four technical (sequencing) replicates of a mouse cecum sample with an initial template input of 1.2 x $10^4$ 16S rRNA gene copies. The red shading indicates the bootstrapped (B = $10^4$) Poisson sampling confidence interval of the input 16S rRNA gene copies. (b) Bootstrapped Poisson sampling relationship between %CV and percentage abundance as a function of read depth.

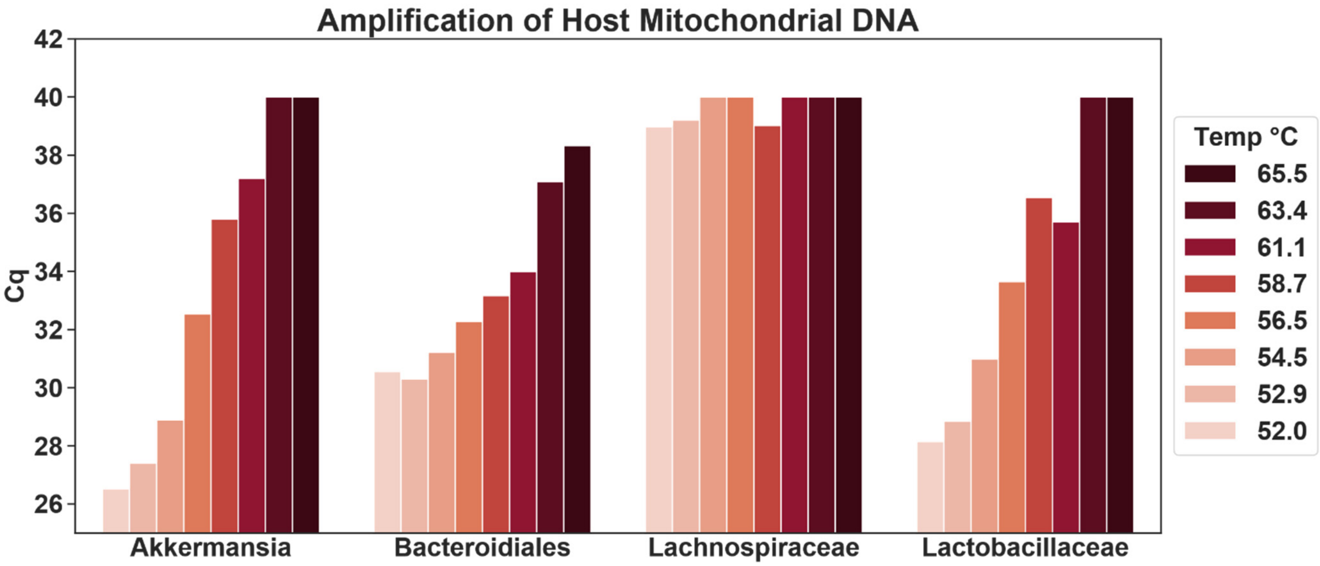


**Figure S5: Optimization of group-specific primers to eliminate amplification of host DNA.** Relative abundance of non-specific product amplified from 20 ng/μL small-intestine mucosa sample from a germ-free mouse measured by qPCR. Lower Cq values indicate more amplification. Each color represents a different annealing temperature used during the cycling process. Samples were run in singlet at each temperature.

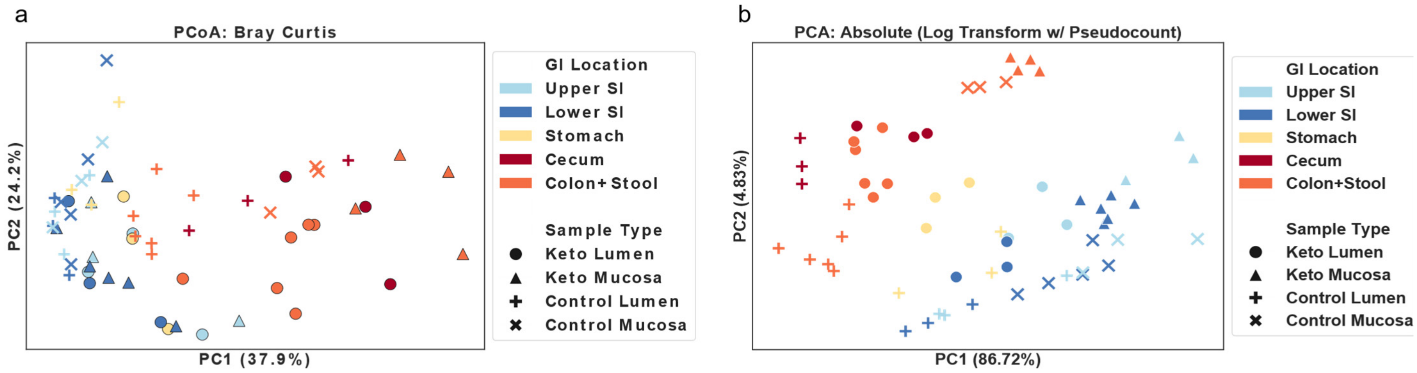


**Figure S6: Impact of ordination method on data visualization**. (a) Principal coordinates analysis (PCoA) plot using Bray–Curtis dissimilarity metric of all samples collected 10 days after the diet switch. (b) Principal component analysis (PCA) plot using log-transform of absolute abundance data after adding a pseudocount of 1 read to all taxa.

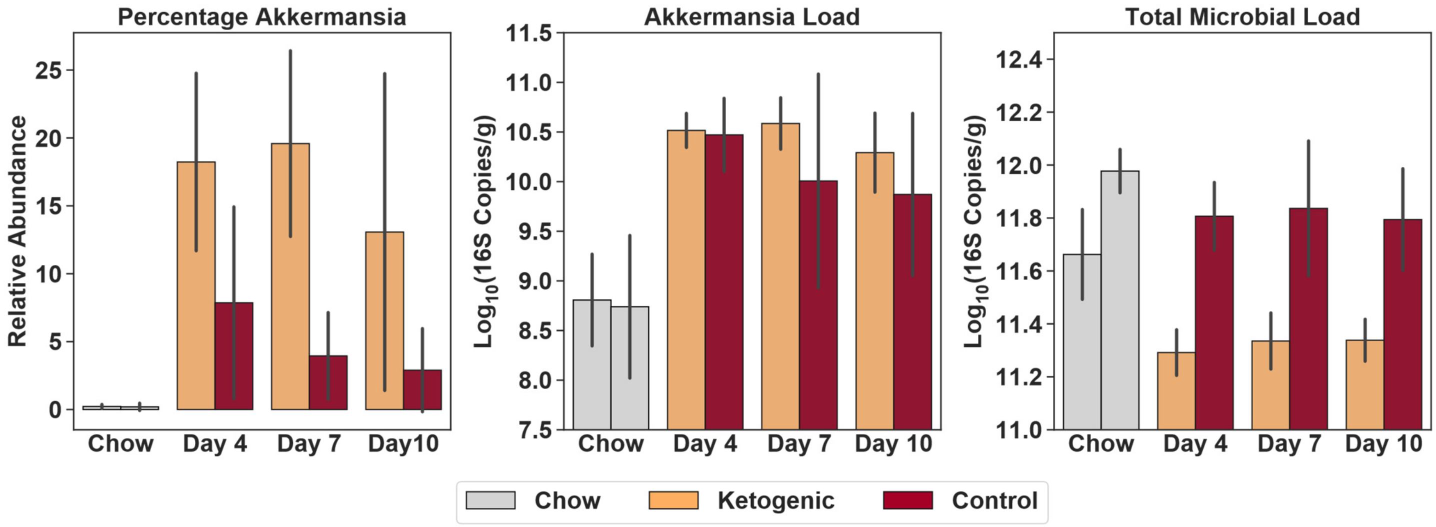


**Figure S7: Comparison of relative and absolute abundance quantification of *Akkermansia(g)* between mice on ketogenic and control diet.** Average *Akkermansia(g)* load from stool of N = 6 mice on control diet and N = 5 mice on ketogenic diet. Bar plots show mean plus or minus the standard deviation.

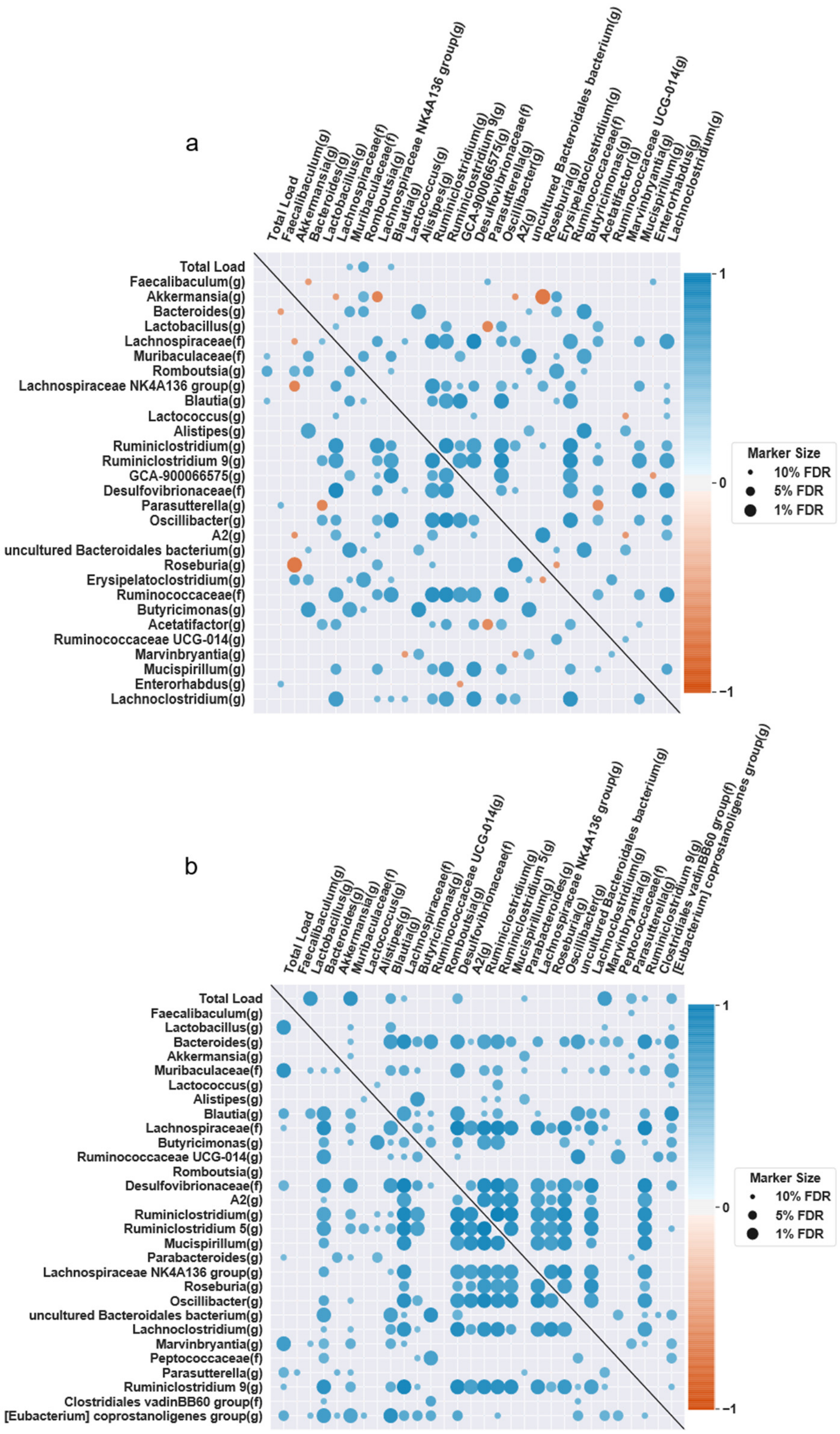


**Figure S8: Absolute-abundance measurements enable unbiased determination of correlation structure in microbiome datasets.** Correlation matrices, using Spearman's rank, for the total microbial load and the top 30 most abundant taxa in stool samples from mice on either a ketogenic diet (a) or control diet (b). The color of each marker is based on the correlation coefficient (orange indicates negative correlations, blue indicates positive correlations) and the size is determined by the q-value of the correlation after Benjamini–Hochberg multiple testing correction. False-discovery rates (FDR) indicate the q-value at which the correlation was deemed significant: 1%, 5%, 10%.

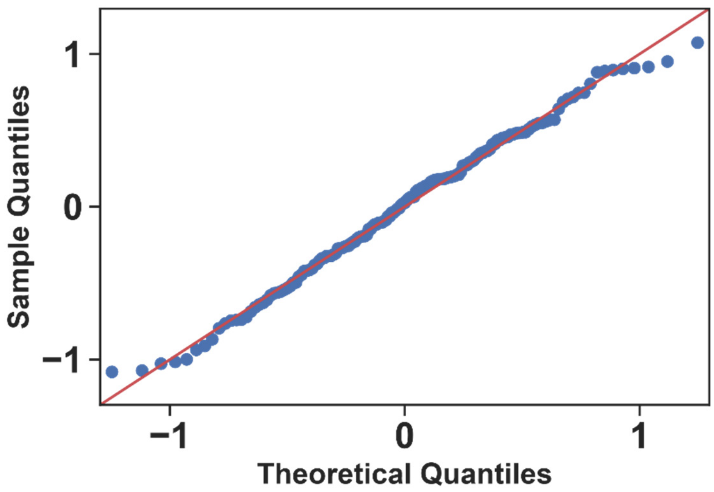


**Figure S9: The uncertainty in taxon absolute-abundance measures approximately follows a normal distribution.** The quantile-quantile (Q–Q) plot of the mean-centered $\log_2$ relative error of absolute taxon abundances. The relative error is calculated as the ratio of the absolute taxon loads measured by our method of quantitative sequencing with dPCR anchoring over the absolute loads measured by taxon-specific primers in dPCR (data are from Fig. 3b). The x-axis represents the theoretical quantiles from a normal distribution while the y-axis is the actual quantiles of the mean-centered $\log_2$ relative errors.

**Table S1: Contaminant taxa with greater than 1% abundance in negative-control extraction.**

| Contaminant Taxa | Percentage Abundance |
|---|---|
| Acinetobacter(g) | 31.38 |
| Pseudomonas(g) | 24.12 |
| Allorhizobium-Neorhizobium-Pararhizobium-Rhizobium(g) | 9.77 |
| Brevundimonas(g) | 5.86 |
| Massilia(g) | 2.84 |
| Delftia(g) | 2.52 |
| Dietzia(g) | 2.33 |
| Corynebacterium 1(g) | 2.08 |
| Xanthomononadaceae(f) | 2.06 |
| Anaerococcus(g) | 1.95 |
| Nubsella(g) | 1.94 |
| Lysobacter(g) | 1.91 |
| Comamonas(g) | 1.82 |
| Janthinobacterium(g) | 1.30 |
| Shinella(g) | 1.29 |
| Novosphingobium(g) | 1.23 |
| Sphingobium(g) | 1.15 |
| Taibaiella(g) | 1.02 |

**Table S2: Comparison between digital PCR anchoring method for absolute abundance measurements and other published absolute abundance methods[2-5].**

| Approach | Major Improvement to the Field | Demonstrated Limit of Quantification | Demonstrated Limit of Detection | Demonstrated Precision | Validated Sampling Locations | Validation Against PCR Amplification Bias | Bias-Free Validation with High Host DNA Loads | Ref. |
|---|---|---|---|---|---|---|---|---|
| **Flow Cytometry** | Showed importance of quantifying absolute abundance in clinical samples | Not Discussed | Not Discussed | Not Discussed | Stool | N/A | Not Shown | Vandeputte *et al.* 2017[2] |
| **Sequencing Spike-ins** | Generated a variety of spike-in standards that can be used. Provided comprehensive analysis of quantitative limits and accuracy | Not Discussed | Dependent on spike-in amount (~100copies/rxn) | 1.5-1.7X, Shown only with mock communities | Sludge, Soil | Show that it can skew total load measurement | Not Shown | Tourlousse *et al.* 2016[3] |
| **qPCR Anchoring** | Provided a simple and easy method for absolute quantification | Not Discussed | Not Discussed | High correlation at high DNA input levels | Stool | Not Discussed | Not Shown | Jian *et al.* 2018[4] |
| **Total DNA** | Provided a simple method for absolute quantification. Showed dramatic variability in loads across animal kingdom and clinical scenarios | Not Discussed | ~100pg of DNA | Not Discussed | Stool | N/A | N/A | Contijoch *et al.* 2019[5] |
| **Digital PCR Anchoring** | Quantitative assessment of accuracy and precision of absolute abundances in complex gut samples and their impact on differential taxon analyses | 4.2x10^5 16S copies/g Stool 1.0x10^7 16S copies/g Mucosa | 4.2x10^4 16S copies/g Stool 1.0x10^6 16S copies/g Mucosa | 2X across 6 orders of magnitude with low and high host DNA load | Stool, Mucosa, Small Intestine, Cecum, Stomach | Yes | Yes | This paper |

**Table S3: Composition of ketogenic and control diets used in this study based on previously reported diets**.[6]

| | TD.150300 | TD.07797.PWD |
|---|---|---|
| | **Control Diet (g/kg)** | **Ketogenic Diet (g/kg)** |
| Casein | 200 | 121 |
| Crisco | 61.25 | 605 |
| Corn Oil | 8.75 | 86.2 |
| Cellulose | 50 | 112.95 |
| Corn Starch | 389 | 0 |
| Maltodextrin | 100 | 0 |
| Sucrose | 150 | 0 |
| DL-Methionine | 3 | 1.56 |
| Vitamin Mix, Teklad (40060) | 10 | 17.8 |
| Choline Bitartrate | 0 | 2.5 |
| TBHQ, antioxidant | 0.07 | 0.14 |
| Mineral Mix, Ca-P Deficient (79055) | 13.37 | 23.8 |
| Calcium Phosphate, dibasic | 7.5 | 24.3 |
| Calcium Carbonate | 6.85 | 4.4 |
| Magnesium Oxide | 0.2 | 0.35 |

**Table S4: Absolute abundance, relative abundance, fold change and quantification class for each differentially abundant taxon in the stool 10 days after diet switch.**

| Taxon | Absolute Abundance Ketogenic Diet (16S copies/g) | Absolute Abundance Control Diet (16S copies/g) | $\log_2$ Fold Change (Keto/Control) | Relative Abundance Ketogenic Diet (%) | Relative Abundance Control Diet (%) | Quantification Class |
|---|---|---|---|---|---|---|
| GCA-900066575(g) | 1.94E+09 | 8.71E+07 | 4.00 | 0.799 | 0.014 | Semi-Quant |
| Ruminococcaceae(f) | 2.02E+09 | 2.43E+08 | 2.87 | 0.909 | 0.033 | Semi-Quant |
| Lachnospiraceae NK4A136 group(g) | 5.34E+09 | 8.57E+08 | 2.58 | 2.362 | 0.135 | Quant |
| Acetatifactor(g) | 5.20E+08 | 6.19E+07 | 2.46 | 0.256 | 0.009 | Semi-Quant |
| Lachnospiraceae(f) | 8.29E+09 | 1.71E+09 | 2.25 | 3.708 | 0.226 | Quant |
| Ruminiclostridium 9(g) | 1.91E+09 | 5.01E+08 | 1.84 | 0.863 | 0.059 | Quant |
| Dorea(g) | 7.79E+07 | 0.00E+00 | 1.36 | 0.032 | 0.000 | Presence/Absence |
| Enterorhabdus(g) | 7.55E+08 | 3.51E+08 | 0.99 | 0.349 | 0.052 | Quant |
| [Eubacterium] xylanophilum group(g) | 8.69E+07 | 1.91E+07 | 0.87 | 0.037 | 0.003 | No Quant |
| Peptococcus(g) | 8.83E+07 | 2.29E+07 | 0.79 | 0.040 | 0.004 | Semi-Quant |
| Candidatus Soleaferrea(g) | 5.91E+07 | 6.21E+06 | 0.78 | 0.026 | 0.002 | No Quant |
| Marvinbryantia(g) | 5.67E+08 | 1.35E+09 | -1.26 | 0.226 | 0.218 | Quant |
| Bacteroides(g) | 8.81E+09 | 3.37E+10 | -1.93 | 3.990 | 5.578 | Quant |
| Faecalibaculum(g) | 1.01E+11 | 3.87E+11 | -1.93 | 46.724 | 54.268 | Quant |
| Prevotellaceae UCG-001(g) | 1.50E+07 | 7.87E+07 | -2.12 | 0.006 | 0.015 | No Quant |
| Bifidobacterium(g) | 3.18E+08 | 1.42E+09 | -2.15 | 0.153 | 0.100 | Quant |
| Muribaculaceae(f) | 1.25E+08 | 5.74E+08 | -2.16 | 0.056 | 0.091 | Quant |
| Ruminiclostridium 5(g) | 2.85E+08 | 1.38E+09 | -2.26 | 0.127 | 0.202 | Quant |
| Ruminococcaceae UCG-014(g) | 4.85E+08 | 2.45E+09 | -2.32 | 0.209 | 0.427 | Quant |
| Ruminococcaceae NK4A214 group(g) | 8.66E+06 | 6.64E+07 | -2.36 | 0.003 | 0.009 | No Quant |
| Lactococcus(g) | 3.34E+09 | 1.74E+10 | -2.38 | 1.528 | 2.715 | Quant |
| Muribaculaceae(f) | 8.04E+09 | 4.26E+10 | -2.40 | 3.520 | 6.506 | Quant |
| Anaerotruncus(g) | 2.23E+07 | 1.79E+08 | -2.68 | 0.010 | 0.031 | No Quant |
| Lactobacillus(g) | 1.45E+10 | 1.35E+11 | -3.22 | 6.632 | 19.295 | Quant |
| Butyricimonas(g) | 4.38E+08 | 4.39E+09 | -3.30 | 0.203 | 0.755 | Quant |
| Alistipes(g) | 1.11E+09 | 1.15E+10 | -3.36 | 0.526 | 2.018 | Quant |
| Mollicutes RF39(o) | 3.06E+07 | 3.99E+08 | -3.38 | 0.014 | 0.074 | Semi-Quant |
| Christensenellaceae(f) | 2.35E+07 | 3.69E+08 | -3.55 | 0.009 | 0.057 | Semi-Quant |
| Clostridiales vadinBB60 group(f) | 3.31E+07 | 5.68E+08 | -3.77 | 0.017 | 0.108 | Semi-Quant |
| ASF356(g) | 0.00E+00 | 2.44E+08 | -4.65 | 0.000 | 0.029 | Presence/Absence |
| Parabacteroides(g) | 9.26E+07 | 3.30E+09 | -5.01 | 0.040 | 0.457 | Quant |
| Gram-negative bacterium cTPY-13(g) | 0.00E+00 | 3.71E+08 | -5.20 | 0.000 | 0.050 | Presence/Absence |

**Table S5: Absolute abundance, relative abundance, fold change, and quantification class for each differentially abundant taxon in the lower small-intestine mucosa 10 days after diet switch.**

| Taxon | Absolute Abundance Ketogenic Diet (16S copies/g) | Absolute Abundance Control Diet (16S copies/g) | $\log_2$ Fold Change (Keto/Control) | Relative Abundance Ketogenic Diet (%) | Relative Abundance Control Diet (%) | Quantification Class |
|---|---|---|---|---|---|---|
| Lachnoclostridium(g) | 1.64E+07 | 7.07E+05 | 3.57 | 0.293 | 0.006 | Semi-Quant |
| Lachnospiraceae(f) | 9.60E+06 | 4.05E+05 | 3.16 | 0.171 | 0.006 | Semi-Quant |
| A2(g) | 2.29E+07 | 2.08E+06 | 3.06 | 0.441 | 0.025 | Semi-Quant |
| Akkermansia(g) | 2.57E+08 | 3.77E+07 | 2.74 | 5.576 | 0.419 | Quant |
| Escherichia-Shigella(g) | 2.96E+06 | 0.00E+00 | 2.21 | 0.059 | 0.000 | Presence/Absence |
| Dorea(g) | 2.94E+06 | 0.00E+00 | 2.20 | 0.062 | 0.000 | Presence/Absence |
| Bacteroides(g) | 5.51E+06 | 8.09E+05 | 1.94 | 0.112 | 0.009 | Semi-Quant |
| Desulfovibrionaceae(f) | 4.29E+06 | 5.91E+05 | 1.82 | 0.097 | 0.005 | Semi-Quant |
| uncultured Bacteroidales bacterium(g) | 1.55E+07 | 3.98E+06 | 1.76 | 0.365 | 0.041 | Quant |
| Enterorhabdus(g) | 1.12E+07 | 2.74E+06 | 1.73 | 0.245 | 0.022 | Semi-Quant |
| Lachnospiraceae NK4A136 group(g) | 1.38E+06 | 3.38E+05 | 0.65 | 0.031 | 0.005 | No Quant |
| Ruminococcaceae(f) | 8.48E+05 | 6.76E+04 | 0.51 | 0.017 | 0.001 | No Quant |
| uncultured Lachnospiraceae bacterium(g) | 6.46E+05 | 0.00E+00 | 0.35 | 0.013 | 0.000 | Presence/Absence |
| Marvinbryantia(g) | 4.62E+06 | 3.33E+06 | 0.27 | 0.127 | 0.037 | Semi-Quant |
| Ruminiclostridium(g) | 5.41E+05 | 0.00E+00 | 0.17 | 0.011 | 0.000 | Presence/Absence |
| Muribaculaceae(f) | 1.22E+08 | 3.48E+08 | -1.51 | 2.585 | 3.041 | Quant |
| Lactococcus(g) | 1.20E+08 | 4.31E+08 | -1.84 | 2.533 | 4.582 | Quant |
| Lactobacillus(g) | 9.05E+08 | 3.70E+09 | -2.03 | 16.956 | 35.988 | Quant |

**Table S6: Primers used in this study, relevant conditions, and specificity**. All primers were tested *in silico* for coverage of their desired taxonomic group and specificity.[7-13]

| | **Akkermansia muciniphila** | **Bacteroidiales** | **Lachnospiraceae** | **Lactobacillaceae** | **519F-806R** |
|---|---|---|---|---|---|
| Forward Primer | CAGCACGTGAAGGTGGGGAC | GGTGTCGGCTTAAGTGCCAT | CGGTACCTGACTAAGAAGC | GCAGCAGTAGGGAATCTTCCA | CAGCMGCCGCGGTAA |
| Reverse Primer | CCTTGCGGTTGGCTTCAGAT | CGGAYGTAAGGGCCGTGC | AGTTTYATTCTTGCGAACG | CACCGCTACACATGGAG | GGACTACHVGGGTWTCTAAT |
| Taxonomy Level | Species | Order | Family | Family | Kingdom |
| Annealing Temp (°C) | 65 | 65 | 55 | 60 | 52 |
| Concentration (nM) | 500 | 500 | 500 | 500 | 500 |
| Coverage (n=1 mismatch) | 100% | 75% | 86% | 91% | 94% Bacteria, 95% Archaea |
| Potential Undetected Taxa | None | Rikenellaceae(f); Alistipes(g) | UCG-010(g) | None | None |
| Potential non-specific interactions (n=1 mismatch) | None | None | None | Leuconostocaceae(o) | None |
| Citation | w | Rinttila *et al.* (2004)[9] | Kennedy *et al.* (2014)[10] | Castillo *et al.* (2006)[11] | Bogatyrev & Ismagilov (2020)[14]<br>Bogatyrev *et al.* (2020)[13]<br>Bogatyrev (2020)[12] |
